# Depth structures metabolic potential and warming responses in marine microbial eukaryotes

**DOI:** 10.64898/2026.11.10.765571

**Authors:** Mattia Greco, Lidia Montiel, Francisco Latorre, Caterina R. Giner, Carlos M. Duarte, Susana Agusti, Silvia G. Acinas, Raffaele Bernardello, Ramon Massana, Ramiro Logares

## Abstract

Microbial eukaryotes are key drivers of food-web dynamics and nutrient cycling across the ocean’s water column. Yet, how their metabolic potential is structured through depth remains largely unknown, limiting our ability to understand how microbial communities mediate biogeochemical transformations and how these functions may shift under climate change. Here, we combined a global-scale metagenomic analysis of tropical and subtropical samples spanning the full water column with bathypelagic metatranscriptomics to identify depth-dependent functional patterns across plankton size fractions. At depth, the functional composition of the nanoplankton size fraction was consistent with the export of surface-derived biomass, whereas the picoplankton size fraction showed signatures of resident communities, including depth-specific heterotrophic functions and bathypelagic-restricted fungal assemblages. Metabolic functional diversity was tightly coupled to taxonomic composition along the water column. Heterotrophic lineages contributed to phosphorus and sulfur acquisition at depth, revealing an undercharacterised eukaryotic role in deep-ocean nutrient cycling. Projections under end-of-century warming revealed spatially structured metabolic responses: subtropical intensification of photosynthetic capacity, subpolar stimulation of aerobic metabolism, and a decline of heterotrophic potential consistent with metabolic rebalancing within mixotrophic taxa. Our findings highlight the importance of marine microbial eukaryotes in ocean biogeochemistry, reveal their distinct functional roles across depths, and provide novel projections of how their metabolic potential may change under ocean warming.

## INTRODUCTION

The vertical structure of the ocean is a fundamental determinant of marine ecosystem functioning (1, 2). From the sunlit epipelagic zone where photosynthesis fuels primary production to the dark abyss where organic carbon is produced only chemosynthetically, respired and remineralised, depth-related gradients in light, temperature, pressure, nutrients, and organic matter create distinct ecological habitats that shape biodiversity and regulate key biogeochemical processes, including nutrient cycling and carbon sequestration. Central to these processes are microbial eukaryotes (protists and fungi; hereafter ME), which drive food-web dynamics and mediate nutrient and carbon cycling (3, 4). These organisms harbor immense diversity, accounting for ∼35 % of the marine biomass (5) and encompassing a wide range of trophic roles, metabolic strategies, and functional capabilities (6–8). Unlike prokaryotes, where horizontal gene transfer frequently decouples taxonomy from functional potential, promoting functional redundancy (9, 10), ME functional roles are deeply constrained by cell size, trophic mode, and morphology, traits that are phylogenetically conserved and rarely transferable across lineages (11). As a consequence, taxonomy and function are more tightly coupled in ME than in prokaryotes (9, 10, 12), so environmentally driven community turnover, whether along depth gradients or under changing ocean conditions, is likely to carry more direct functional consequences.

Multiple studies have demonstrated that ME community composition and interactions shift markedly across depth gradients (13–19). However, our understanding of how depth-driven changes in microbial eukaryotic diversity translate into shifts in community functional repertoires, and how these functional changes ultimately influence key ecosystem processes, remains limited (20, 21). Eukaryotic metatranscriptomics has begun to characterise functional biogeography across size fractions within the euphotic zone (22), and depth-resolved analyses of prokaryotic communities have revealed that community turnover is the dominant driver of gene expression variation with depth (23). Yet, a comparable depth-resolved functional characterisation of ME across size fractions is still lacking. Addressing this gap is particularly urgent as climate change reshapes vertical ocean structure, driving cascading changes in microbial communities and ecosystem function (24).

Predicting how microbial communities respond functionally to climate change is recognised as a major challenge (25). However, research has focused almost entirely on prokaryotic and phytoplankton groups (25–27), leaving projections of ME metabolic potential largely unexplored. Given the taxonomy–function coupling, functional projections for ME would be a particularly valuable component of ocean climate prediction (28).

Recent advances in marine metagenomics provide a powerful framework for investigating both the depth-structuring of ME functional potential and its environmental drivers (21). Leveraging this approach, we analysed 116 marine metagenomes (MetaG) and from 11 vertical profiles collected during the circumglobal Malaspina Expedition, spanning the epipelagic (0–200 m), mesopelagic (200– 1,000 m), and bathypelagic (>1,000 m) zones, complemented by 51 bathypelagic metatranscriptomes (MetaT) (29) (Fig. 1). We developed a novel bioinformatic framework to improve the identification of eukaryotic genes from metagenomes and minimize prokaryotic contamination, enabling the recovery of an eukaryotic gene catalogue comprising ∼7 million genes (or Open Reading Frames; ORFs). We quantified how pico-eukaryotes (0.2–3.0 μm cell-size) and nano-eukaryotes (3.0–20 μm) differentially contribute to community turnover and gene abundance patterns across depths for key biogeochemical processes, including photosynthesis and the cycling of carbon, nitrogen, phosphorus, and sulfur. We then used bathypelagic metatranscriptomes to assess whether depth-structured functional potential is reflected in gene expression in the dark ocean. Next, by directly comparing the functional profiles of eukaryotic and prokaryotic organisms, we reveal previously uncharacterized contrasts in their depth-resolved metabolic organisation. Finally, we identified robust associations between metabolic pathways and environmental variables and used these relationships to project how depth-structured eukaryotic metabolic potential may respond to end-of-century ocean conditions under the SSP5–8.5 scenario.

**Fig. 1.**
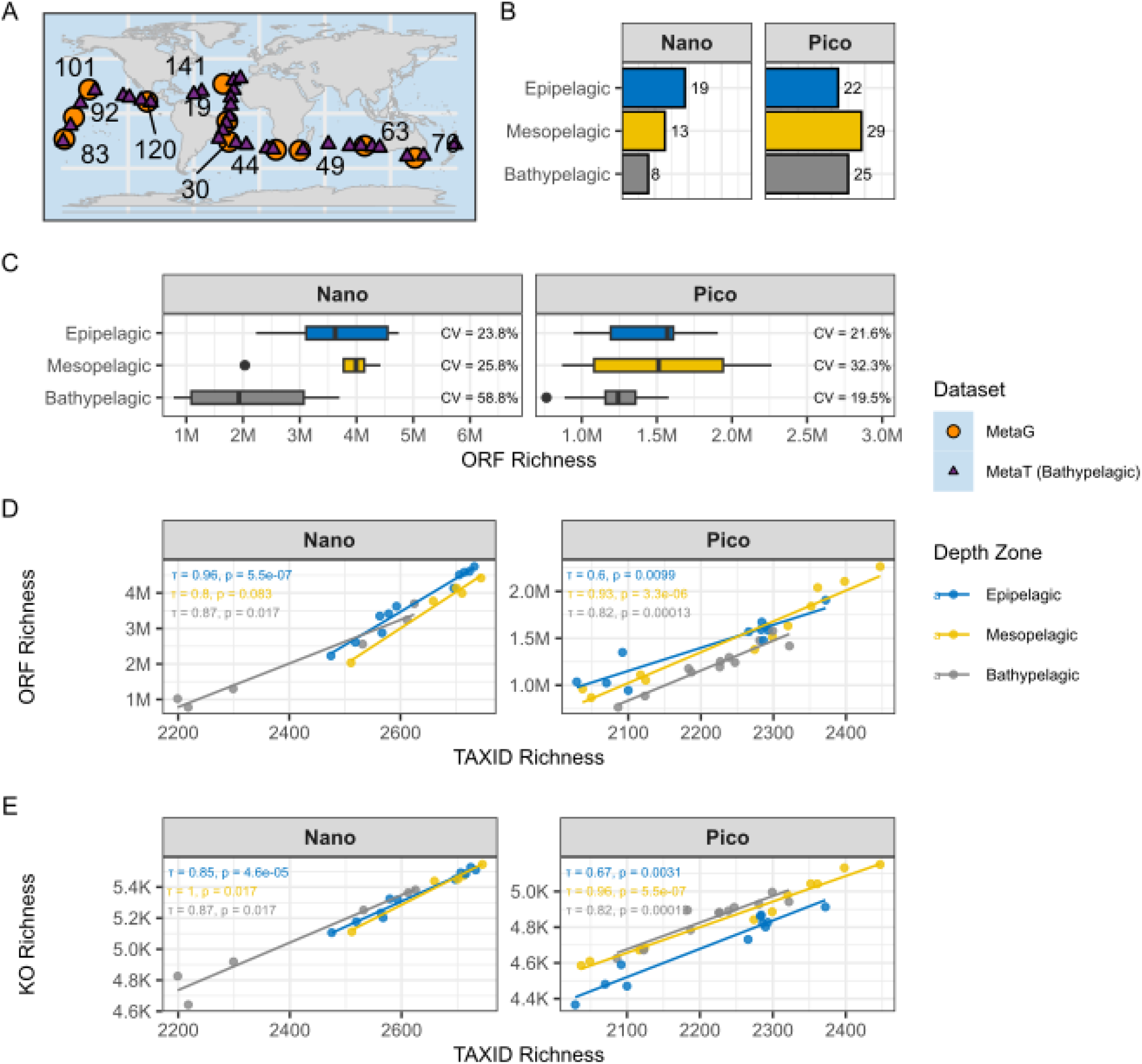
The MALASPINA marine microbial eukaryote gene catalogue across depth zones. (A) Sampling stations from the Malaspina expedition, showing metagenomic (MetaG) and bathypelagic metatranscriptomic (MetaT) locations (Station IDs shown only for MetaG). (B) Number of samples (MetaG) per depth zone for nano and pico size fractions. (C) MetaG-derived ORF richness per sample across depth zones for nano (top) and pico (bottom) fractions; the coefficient of variation (CV) is indicated for each depth zone. (D) Correlation between TAXID richness and ORF richness for nano (top) and pico (bottom) fractions, coloured by depth zone. (E) Correlation between TAXID richness and KO richness for nano (top) and pico (bottom) fractions, coloured by depth zone. Kendall’s τ and associated p-values are shown in panels D and E.

## RESULTS

### Coupling Between Taxonomic and Functional Richness Across Depth Layers

The curated microbial eukaryote catalogue comprised 7,030,775 ORFs, with over half assigned to Haptophyta (27.7%), Dinophyceae (17%), and Coscinodiscophyceae (6%) (Fig. S1). In both size fractions, phototrophic taxa were consistently detected even in deeper layers, including the bathypelagic zone. The catalogue was compared per taxonomic group to 18S rDNA mTags from the same metagenomes (Table S14, Fig. S1). Phototrophs dominated the ORF catalogue but were a minor fraction of mTag reads at every depth. The amplicon-defined MALV and RAD environmental clades reached 17–23% of epipelagic mTag reads but were absent from the catalogue. Dinoflagellates were more abundant in mTags than ORFs, whereas diatoms and other well-referenced phototrophs showed the opposite pattern (See Limitations).

In terms of functional richness, the nano fraction contained the largest number of ORFs overall (Fig. 1C), and accumulation curves showed consistently higher richness and earlier saturation in nano than pico fractions for both ORFs and KOs (Fig. S2). The same analysis also suggests that additional sampling (particularly in the pico fraction) would continue to recover new ORFs as functional diversity approaches saturation.

For the nano fraction, the epipelagic layer exhibited the highest richness, with a maximum of 4,743,977 ORFs recorded at station 63 (Indian Ocean) (Fig. 1C). ORF richness declined by 43% from the epipelagic to the bathypelagic layer, reaching its lowest value in the bathypelagic zone (778,001 ORFs at station 92, South Pacific), which also displayed the greatest variability among samples (CV = 59%) (Fig. 1C). In contrast, within the pico fraction, the highest richness was observed in the mesopelagic layer (Fig. 1C), with a maximum of 2,263,386 ORFs at station 141 (North Atlantic), which also exhibited the greatest variability in richness across stations (CV = 32%). Similar to the nano fraction, richness in the pico community decreased with depth, but the decline was less pronounced (20%). The lowest richness in the pico fraction occurred in the bathypelagic zone (765,215 ORFs at station 44, Indian Ocean); however, unlike the nano fraction, this depth layer showed the lowest variability among samples (CV = 19.5%).

Taxonomic and functional diversity were strongly correlated across depth zones in both the pico and nano microeukaryotic communities (Fig. 1D and 1E). Consistent with the ORF-based annotations, a greater number of KEGG KOs were identified in the nano fraction compared with the pico fraction (Fig. 1E). All correlations were highly significant. The relationship between taxonomic and functional richness was strongest in the epipelagic layer for the nano community, whereas in the pico community it peaked in the mesopelagic and weakened toward the surface (τ = 0.64; Figs. 1D-E).

To examine depth-associated differentiation across taxonomic and functional levels, we identified depth-specific distinctive features (i.e., features absent from other depth zones), hereafter referred to as “distinctiveness,” at the Class (taxonomic), ORF, and KEGG Orthology (KO) levels (Fig. 2). Distinctiveness patterns differed markedly between size fractions.

**Fig. 2.**
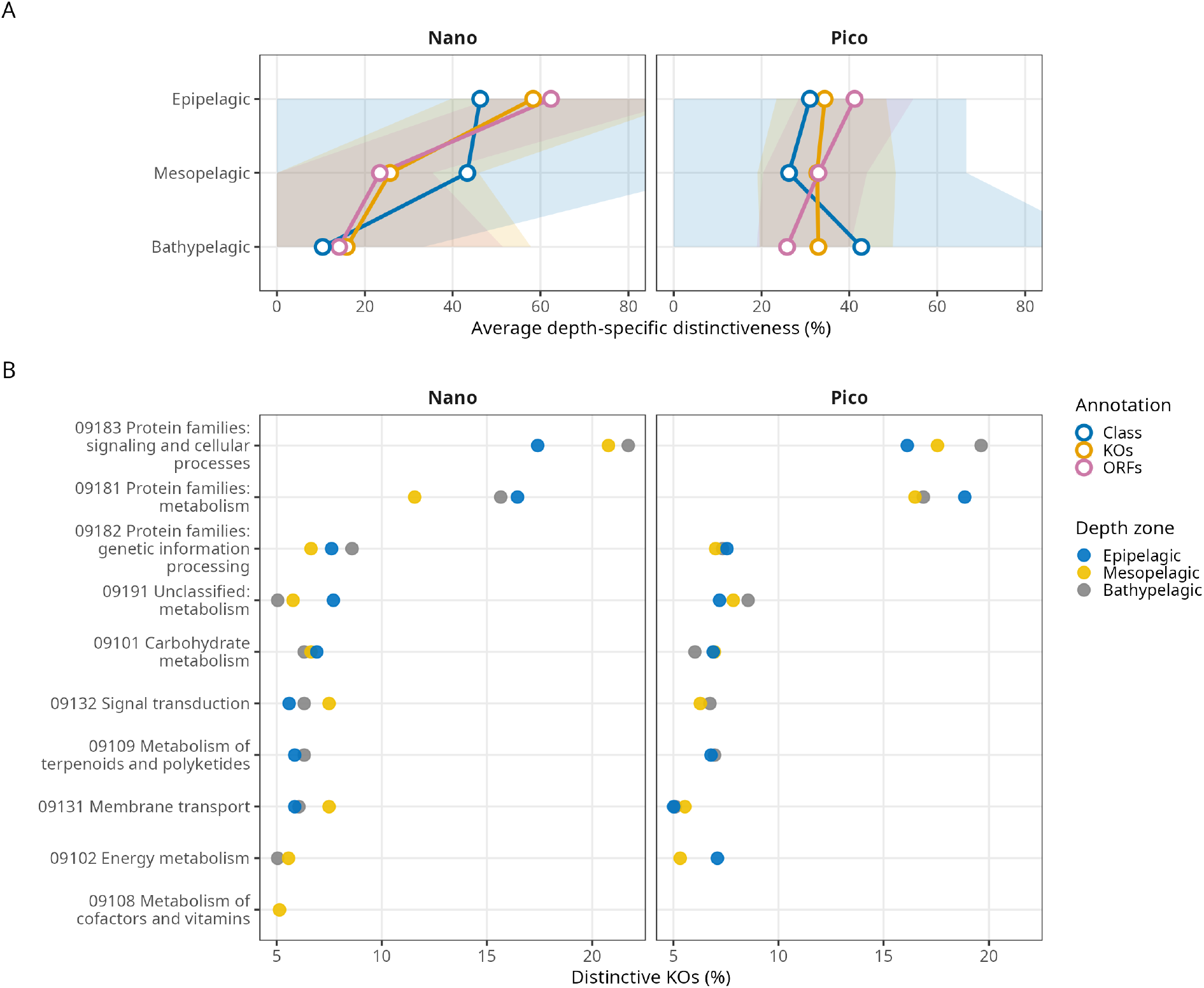
Depth-exclusive functional and taxonomic features across size fractions. (A) Average depth-specific distinctiveness (%) across epipelagic, mesopelagic, and bathypelagic zones for three annotation types (taxonomic class, KOs, and ORFs), shown separately for nano (3–20 µm) and pico (0.2–3 µm) size fractions. Shaded ribbons represent the observed range (minimum– maximum) across stations. (B) Percentage of depth-distinctive KOs assigned to each KEGG BRITE functional category, shown per depth zone (colour) and size fraction. Only categories exceeding 5% of distinctive KOs in at least one depth zone are shown.

In the nano fraction, all three annotation levels showed high depth-specific distinctiveness in the epipelagic, with functional markers (KOs and ORFs) reaching ∼60% and taxonomic classes ∼40% (Fig. 2A). The transition to the mesopelagic was marked by a sharp collapse in functional distinctiveness while taxonomic distinctiveness remained comparably high. From mesopelagic to bathypelagic, all annotation levels declined to similarly low values. This pattern indicates that each depth zone harbours a taxonomically distinctive community, but that only the epipelagic community translates taxonomic distinctiveness into a correspondingly distinctive functional repertoire, as below the photic zone, taxonomically distinct assemblages converge on similar functional profiles.

In contrast, the pico fraction showed a more complex pattern. Functional distinctiveness at the KO level was remarkably stable across all depth zones, while ORF-level distinctiveness declined modestly from the epipelagic to the bathypelagic (Fig. 2A). Taxonomic distinctiveness followed a different trajectory: a slight decrease from epipelagic to mesopelagic, followed by a sharp increase in the bathypelagic, driven largely by the emergence of exclusive fungal classes. Thus, depth-driven differentiation in pico communities was primarily taxonomic in the bathypelagic, while functional repertoires (particularly at the KO level) remained relatively consistent across the water column.

Despite these contrasting depth profiles, annotation of distinctive KOs into BRITE functional categories revealed substantial overlap between fractions, particularly in the mesopelagic and bathypelagic. There, protein family metabolism and signalling pathways dominated the distinctive signal in both nano and pico (Fig. 2B), suggesting that the broader functional categories showing depth differentiation are generally conserved across size fractions.

### Marker-resolved metabolic stratification across size fractions

Metabolic marker enrichment revealed pronounced size and depth-dependent functional structuring among microbial eukaryotes. Pico-eukaryotes displayed a greater number of enriched markers overall (Fig. S3), indicating broader functional versatility relative to nano-eukaryotes. Across both Prokaryotic and Eukaryotic domains, carbon metabolism and phototrophy markers were, as expected, most strongly enriched in epipelagic waters, although a subset persisted at depth (Fig. 3).

**Fig. 3.**
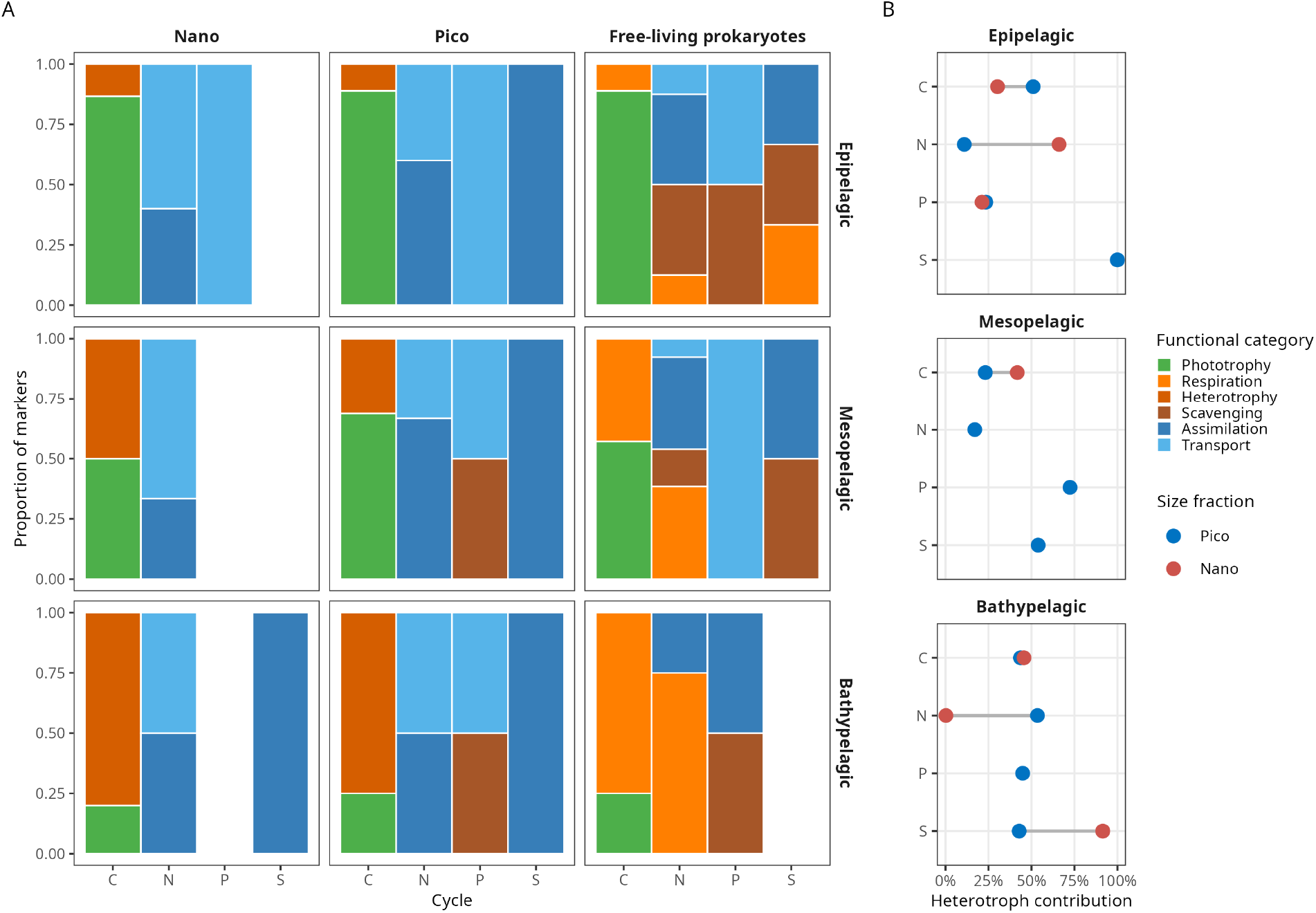
Functional composition of significantly depth-enriched markers in nano- and pico-eukaryotic size fractions and free-living prokaryotes. (A). Columns represent nutrient cycles (C, carbon; N, nitrogen; P, phosphorus; S, sulfur); bar height reflects the relative proportion of each functional category (Phototrophy, Heterotrophy, Assimilation, Respiration, Transport, Scavenging). Rows correspond to depth zones (Epipelagic, Mesopelagic, Bathypelagic). Dumbbell plot (B) shows the contribution of putative heterotrophic taxa to enriched markers per nutrient cycle and depth zone, comparing nano (red) and pico (blue) fractions; values represent the proportion of mean relative abundance attributable to heterotrophic eukaryotic groups.

Heterotrophic potential diverged between size fractions. In nano-eukaryotes, markers such as *SEC61A* (phagosome) and fatty acid degradation genes (*HADHB/ACOX*) were enriched in surface waters, whereas pico-eukaryotes showed enrichment of heterotrophy-related markers predominantly in mesopelagic and bathypelagic layers, indicating vertical partitioning of trophic strategies. Functions from free-living prokaryotes exhibited a parallel shift, with respiratory genes including *coxL*, *ccoN*, and *cydA* enriched at depth.

Nitrogen metabolism displayed clear depth stratification. In surface waters, pico-eukaryotes were associated with ammonium uptake and assimilation (e.g., *MEP/AMT*, *GLN1/GLUL/GS*, *nirA*), while nano-eukaryotes exhibited enrichment of both ammonium and nitrate transporters ( *MEP/AMT*, *NRT2*). At depth, pico-eukaryotes showed broader enrichment of nitrogen acquisition pathways (*NRT*, *nasB*, *nasC/nasA*, *nirB*, *narH*, *GLN1/GLUL/GS*), whereas nano-eukaryotes were enriched primarily in nitrate-associated markers (*narB*, *narH*, *NRT*). Functions from free-living prokaryotic communities similarly displayed enrichment of nitrate respiration and assimilation genes (*narG*, *napA*, *nirB*, *nifH*) in deeper waters.

Phosphorus metabolism showed strong size-dependent differentiation. In the nano fraction, phosphorus cycling markers were restricted to the epipelagic, where transport genes (*PHO84*) were contributed predominantly by Haptophyta, Dinophyceae, Dictyochophyceae, and Bacillariophyta, with no meaningful signal at mesopelagic or bathypelagic depths (Fig. S6). In the pico fraction, phosphorus transport markers were detected across all depth zones and remained dominated by photosynthetic taxa. Phosphorus scavenging (*phoD*) and transport markers (*TC.PIT*, *PHO84*) showed a marked depth-dependent shift: absent in the epipelagic, they emerged in the mesopelagic where Bacillariophyta contributed the majority of the signal, before shifting in the bathypelagic toward a community including Chrysophyceae, Fornicata, Foraminifera, Choanoflagellata, and Ascomycota (Fig. S6). Free-living prokaryotic phosphorus cycling genes (*phoA*, *phoD*, *ptxD*) were enriched throughout the water column, particularly in bathypelagic layers.

Sulfur assimilation markers (*SAT*, *APR*, *SiR*) were enriched in both eukaryotic size fractions. In the pico fraction, markers were enriched across epipelagic and mesopelagic zones, with contributions dominated by Acantharea (Fig. S7). In the nano fraction, sulfur assimilation was restricted to the bathypelagic, where *SiR* enrichment was driven predominantly by Choanoflagellata (Fig. S7). In contrast, prokaryotic sulfur cycling genes (*aprA*, *dmdA*) were more prominent in epipelagic and mesopelagic waters.

Bathypelagic-enriched markers of heterotrophy (β-oxidation), nitrogen acquisition, phosphorus scavenging, and sulfur assimilation were actively transcribed at depth, with signal retention across stringent mapping-identity thresholds (75–100%), consistent with eukaryotic expression (Fig. S8). Phototrophy-associated markers were also detected and actively transcribed at depth.

To assess whether the signal from non-picoeukaryotic heterotrophic taxa in the pico size fraction could reflect DNA leakage from the nano or larger size fractions, we compared absolute read counts across size fractions for each heterotrophic group. In all cases, read counts were higher in the pico than in the nano fractions (median pico/nano ratio ranging from ∼1.2 to ∼4.5; Fig. S9), arguing against contamination as the primary explanation for the presence of larger taxa in the pico fraction.

Gene set enrichment analysis revealed distinct patterns of depth invariance and variability across size fractions (Fig. S10). Pathways associated with lower z-scores of the coefficient of variation (CV), indicating greater depth invariance, were predominantly related to core cellular functions in both nano and pico fractions, including genetic information processing, translation, DNA replication and repair, and amino acid metabolism. The nano fraction additionally showed enrichment of central metabolic and biosynthetic pathways, including glycolysis/gluconeogenesis, the citrate (TCA) cycle, and fatty acid biosynthesis, whereas the pico fraction showed a broader representation of pathways related to genetic information processing and cellular machinery, including RNA biogenesis, ribosome structure, and translation factors, alongside selected metabolic functions such as lipoic acid and amino acid metabolism (Fig. S10). Together, these patterns indicate strong conservation of fundamental cellular processes across depth layers in both size fractions. In contrast, pathways associated with higher CV z-scores, indicating greater depth variability, were consistently related to photosynthesis, including photosystems and antenna proteins, as well as membrane transport and carbohydrate utilization, such as the phosphotransferase system in the nano fraction (Fig. S10).

### Environmental and depth gradients shape functional composition

Kendall correlation and GSEA analyses revealed that functional composition in both size fractions was structured along consistent environmental gradients, although the nature of these associations differed between shared and fraction-exclusive pathways (Fig. 4). Because depth, temperature, and inorganic nutrients strongly covary in the open ocean, these correlations do not allow robust attribution of functional associations to any single environmental driver. Nevertheless, they collectively identify pathways characteristic of warmer, nutrient-depleted, sunlit surface waters versus colder, nutrient-rich, dark deep waters.

**Fig. 4.**
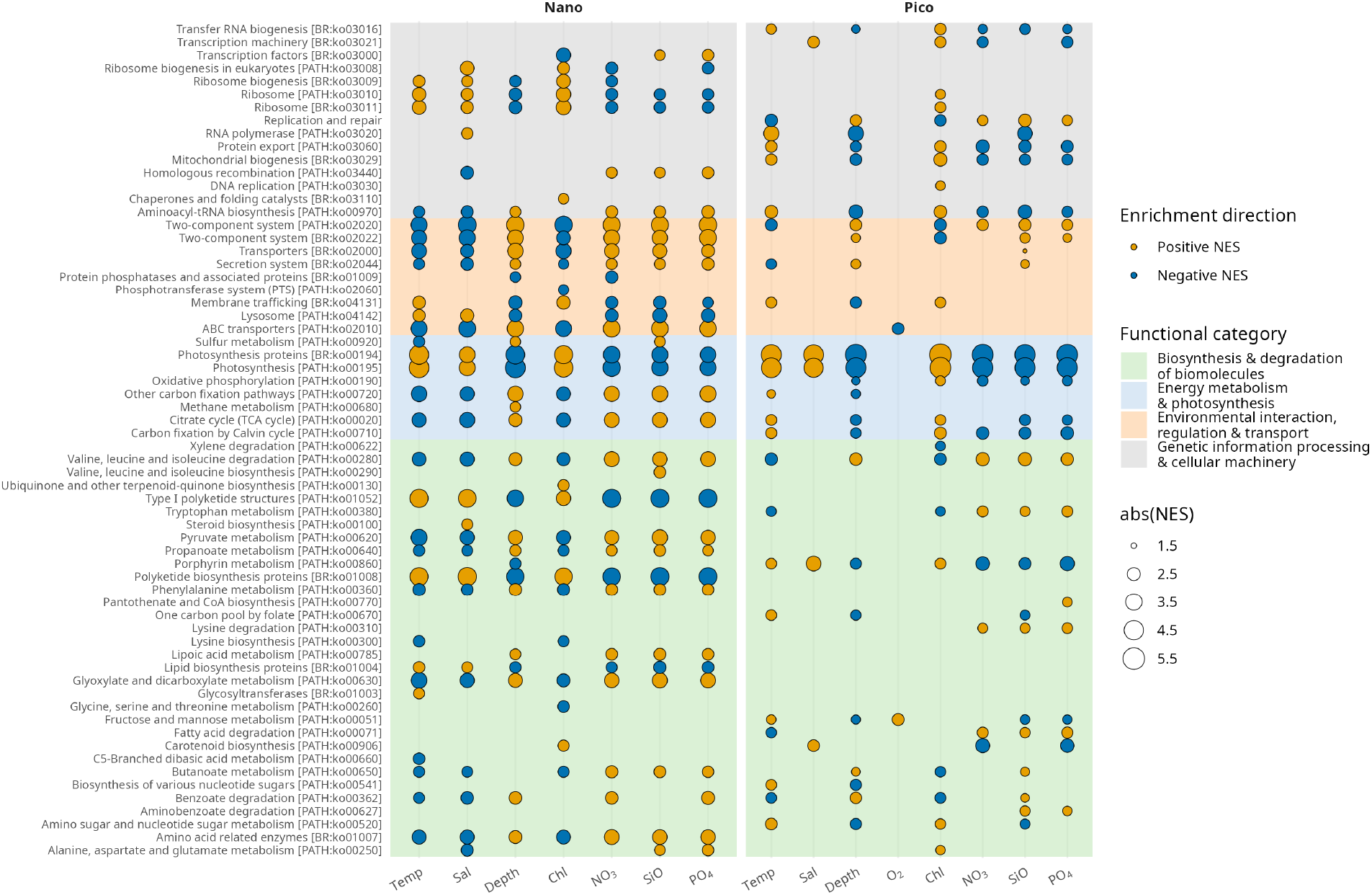
Environmental drivers of microbial eukaryotic functional composition across size fractions. Bubble plot showing GSEA-normalised enrichment scores (NES) for KEGG BRITE pathways against environmental variables, for nano (left) and pico (right) size fractions. Bubble colour indicates enrichment direction (yellow: positive; blue: negative) and bubble size indicates absolute NES magnitude. Pathways are grouped by functional category (colour shading). See Methods for details.

Based on the top pathways ranked by the absolute magnitude of their normalized enrichment scores (NES), GSEA identified 64 unique pathways showing strong associations with environmental gradients. Of these, 43 pathways were unique to one size fraction, whereas 21 were enriched in both fractions. Positive and negative NES values indicated associations with higher and lower values, respectively, of the corresponding environmental variables. Among the shared pathways, 76% showed similar correlation patterns with environmental variables, indicating broadly comparable associations of pico- and nano-sized communities with environmental gradients for this subset of functions. Photosynthesis and carbon fixation pathways (ko00195, ko00194), for example, were positively correlated with chlorophyll, salinity, and temperature and negatively correlated with depth, nitrate, and silicate in both fractions, consistent with their association with surface conditions. Pathways involved in genetic information processing, including RNA polymerase (ko03020) and ribosome (ko03010, ko03011), were also positively correlated with chlorophyll and temperature, particularly in the pico fraction.

In contrast, several pathways involved in transport, signalling, and cellular processes showed opposing environmental associations. ABC transporters (ko02010), for example, were positively correlated with depth, oxygen, and silicate and negatively correlated with temperature, whereas membrane trafficking (ko04131) showed the opposite pattern. Two-component systems (ko02020, ko02022) and secretion systems (ko02044) were negatively correlated with chlorophyll and temperature and positively correlated with depth, nitrate, and silicate, consistent with their association with deeper, nutrient-replete conditions.

Some pathways enriched in both fractions also showed contrasting correlation patterns between size classes. The citrate (TCA) cycle (ko00020) was positively correlated with temperature and negatively correlated with depth in the pico fraction, but showed the opposite pattern in the nano fraction. A similar divergence was observed for aminoacyl-tRNA biosynthesis (ko00970), which was positively correlated with temperature and negatively correlated with depth in the pico fraction, but negatively correlated with temperature and positively correlated with depth in the nano fraction.

Fraction-specific environmental associations were also evident. In the pico fraction, tryptophan metabolism (ko00380), fatty acid degradation (ko00071), protein kinases (ko01001), and replication and repair (ko99976) were positively associated with depth. In the nano fraction, polyketide biosynthesis (ko01008) was associated with surface conditions, whereas sulfur metabolism (ko00920), pyruvate metabolism (ko00620), and glyoxylate and dicarboxylate metabolism (ko00630) were associated with greater depth.

### Projected shifts of microbial eukaryotic metabolic potential across the oceans

We projected end-of-century shifts in the relative abundance of functionally informative KEGG BRITE pathways under the warming scenario SSP5–8.5. Under this scenario sea-surface temperature is projected to increase globally by ∼2.9 °C by 2081–2100 relative to 1995–2014 (30). In the ensemble used here, this corresponds to mean warming of ∼2.2 °C in the epipelagic, ∼1.0 °C in the mesopelagic, and ∼0.1 °C in the bathypelagic between 2015–2020 and 2095–2100. Pathways were selected from those showing significant normalised enrichment scores in the temperature-based GSEA analysis in both size fractions (Fig. 4), prioritising those with consistent temperature associations across size fractions and clear ecological interpretability in the context of microbial eukaryote metabolism. Photosynthesis-related functions capture the primary production capacity of the community and its response to the light and thermal regime of surface waters. Porphyrin metabolism underpins chlorophyll and haem biosynthesis, connecting community metabolic activity to light-harvesting capacity. The citrate cycle (TCA) and aminoacyl-tRNA biosynthesis reflect core biosynthetic and metabolic activity. Membrane transporters integrate nutrient acquisition across trophic strategies. Heterotrophy-related KO markers, representing the community’s catabolic potential, were additionally included as the aggregated abundance of individual KOs that showed significant correlations with temperature in both size fractions, primarily comprising genes involved in fatty acid β-oxidation. Together, these pathways span the major axes of vertical and thermal functional variation in microbial eukaryotic communities, providing an ecologically interpretable window onto the projected restructuring of eukaryotic metabolic potential under warming. Projections were fitted on the pico fraction only, as nano GAMs yielded insufficient explanatory power in temperature-only models (Table S15).

Under the SSP5–8.5 warming scenario, projections indicated pronounced and depth-dependent changes in the relative abundance of BRITE pathways and functional markers in picoeukaryote communities (Fig. 5). No significant change was projected for any category in the bathypelagic zone, which is therefore not shown.

**Fig. 5.**
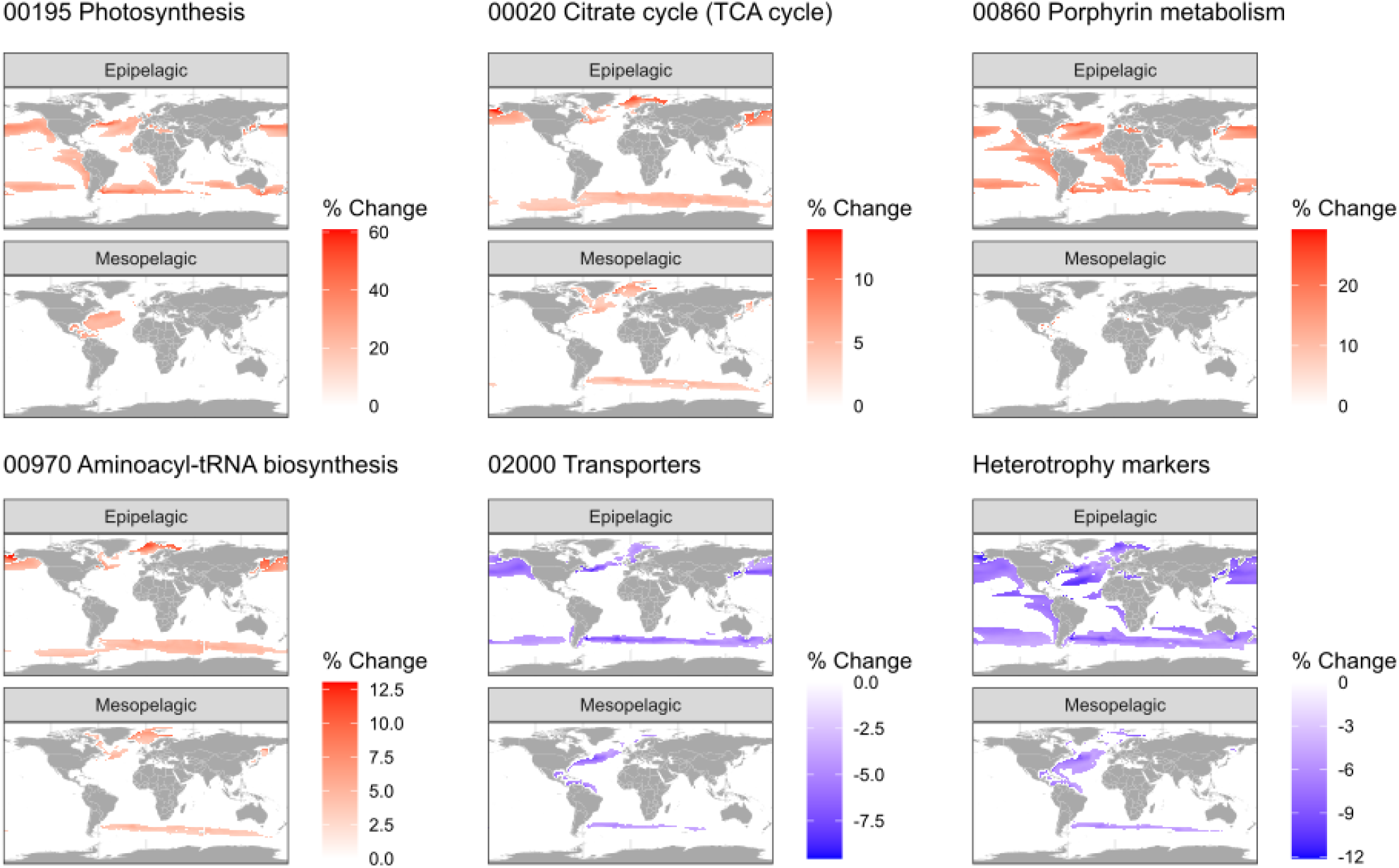
Projected changes in picoeukaryote functional potential under end-of-century ocean warming. Projected percentage change in relative abundance of six functional categories in the pico size fraction in the epipelagic and mesopelagic zones, between 2015–2020 and 2095–2100 under SSP5-8.5 (mean projected warming: ∼2.2 °C epipelagic, ∼1.0 °C mesopelagic).The bathypelagic zone is not shown, as projected warming there is minimal (∼0.1 °C) and no pixel showed a significant change (|Δ| < 1.96 SE everywhere).

Photosynthesis-related pathways exhibited the strongest response, with increases of ∼15–60% in the epipelagic layer, peaking (>40%) at around 40° latitude in both hemispheres, and of ∼20–40% in the mesopelagic western North Atlantic. Similarly, the citrate (TCA) cycle showed moderate increases (3– 14%), most prominently in epipelagic subpolar waters; in the mesopelagic of the western North Atlantic, north of Iceland, the Fram Strait, the Barents Sea and austral subpolar waters. Porphyrin metabolism increased by approximately ∼11–18% (locally up to ∼29%) in the epipelagic layer of tropical and subtropical regions but remained mostly unchanged in deeper layers. Aminoacyl-tRNA biosynthesis also increased in surface and mid-depth waters, with epipelagic increases of ∼3–13% in subpolar regions and mesopelagic increases of ∼3–6% (up to ∼10% in the Nordic and Barents Seas). Derivative-based testing found no statistically significant reversal in any fitted temperature response within the sampled range (0–29°C): two categories (transporters, heterotrophy) declined monotonically across the full range, while apparent non-monotonicity in the point-estimate curves for porphyrin metabolism and aminoacyl-tRNA biosynthesis was not distinguishable from a plateau and did not reflect a confirmed reversal (Table S7). In contrast, transport-related functions declined in both the epipelagic (3–10% reduction) and mesopelagic layers, particularly in the North Pacific, northwest Atlantic, Barents Sea, and austral subpolar waters. Mesopelagic declines (∼3–8%) were concentrated in the North Atlantic and the Barents Sea. Markers associated with heterotrophy also declined, with reductions of up to 12% in epipelagic waters of the North Atlantic, North Pacific, and Southern Ocean, and similar negative trends in the mesopelagic (∼4–10%) of the northwest and South Atlantic.

Single-predictor models based on NO₃ and Si explained a similar share of variance to the temperature models (R^2^ = 0.38–0.86; Table S12). Nutrient-driven projections were spatially coherent with each other (r = 0.70–0.88 across depth zones) but only weakly aligned with temperature-driven projections (r = −0.06 to 0.42 across temperature products; Table S16). This decoupling was consistent with changes in the projected climate fields themselves. NO₃ and Si remained tightly correlated from the training period into the projections (0.81 → 0.88–0.97), whereas the strong temperature–nutrient correlation in the training data weakened in the projections, modestly in the epipelagic and substantially at depth (Table S10). Nevertheless, where projections were significant, all three predictors agreed on the direction of change for five of six functional categories, with >90% sign concordance in the epipelagic (Fig. S11). This result was robust to reduced model flexibility (Table S16), and significant projections fell almost entirely within the core of the training data (Table S8). The exception was aminoacyl-tRNA biosynthesis, for which no significant Si-driven change was detected.

## DISCUSSION

### Depth structuring of taxonomic and functional diversity

Haptophyta and Dinophyceae were the largest contributors to the gene catalogue, extending their well-documented dominance in ribosomal gene surveys (31, 32) to functional gene content, concordant with a western North Atlantic metatranscriptome survey (33). Their contribution remained high throughout the water column, although this pattern partly reflects the availability of reference genomes. In the same samples, 18S data show both groups declining with depth as heterotrophic and parasitic clades become dominant (see Limitations).

Photosynthesis was the primary axis of vertical functional change, with photosynthesis-related pathways showing the highest NES in both size fractions (Fig. S10). Core cellular machinery, including ribosomal, translational, and central metabolic functions, varied little across depth zones, consistent with their constitutive housekeeping roles.

The photosynthetic signal nonetheless persisted into the bathypelagic, where photosystem and Calvin-cycle markers were among the most highly transcribed eukaryotic genes (Fig. S8). This may reflect the persistence of intact phototrophs during sinking, with some cells remaining viable and metabolically active as they are exported to depth (14, 34–37).

The two size fractions differed in how diversity was structured with depth. Nanoeukaryotic ORF richness declined steeply with depth (Fig. 1C), and their functional distinctiveness was concentrated in the epipelagic. This pattern is consistent with a community more strongly associated with photic-zone conditions and with its representation at depth being linked to sinking and export. Picoeukaryotes, by contrast, showed stable functional distinctiveness across depth zones, consistent with more depth-specific resident communities. Their ORF richness peaked in the mesopelagic, extending an amplicon-based pattern from the same samples (14). This maximum was consistently associated with stations harbouring oxygen minimum zones (OMZ) and/or deep scattering layers (DSL) (stations 83 and 141). OMZs sustain diverse picoeukaryotic assemblages through high bacterivory by mixotrophic and heterotrophic taxa (14), while DSLs deliver daily pulses of labile organic matter via diel vertical migration of small fish, fuelling mesopelagic microbial diversity and metabolism (38). Together, these features may create diversity and activity hotspots that elevate picoeukaryotic functional richness in the mesopelagic. Below the photic zone, depth-specific KOs in both fractions converged on protein family metabolism and signalling, suggesting a shared functional response to deep-ocean conditions.

Taxonomic and functional richness were strongly correlated across depth zones and size fractions, and this coupling persisted into the bathypelagic. This suggests that taxonomic diversity can provide a useful proxy for functional diversity in marine microbial eukaryotes, contrasting with the stronger functional redundancy reported in many prokaryotic systems (9, 10). This pattern may reflect the more lineage-specific functional roles of eukaryotes, whose ecological functions depend strongly on morphological and behavioural traits (11, 12, 39). Coupling in richness did not, however, extend to compositional turnover. In the bathypelagic pico fraction, taxonomic distinctiveness increased, driven largely by putatively fungal classes (Dimargaritomycetes, Zoopagomycetes, Archaeorhizomycetes, Olpidiomycetes), whereas KO distinctiveness remained stable. This could indicate that these lineages contribute functions already represented in the broader community, pointing to functional redundancy (39), although incomplete KO annotation could produce the same pattern. The fungal assignments warrant caution given the paucity of marine fungal references (40). Nevertheless, their consistent restriction to bathypelagic depths across stations supports the biological relevance of the pattern. This agrees with reports of abundant bathypelagic Basidiomycota (41), fungal dominance of deep marine snow (42) and evidence from the same Malaspina samples that these are metabolically active yeasts whose abundance tracks refractory carbon (43). Together, these findings are consistent with resident deep-sea fungal communities rather than passive sinking alone (44).

### Depth-dependent differentiation of metabolic functions

A clear size-fraction niche differentiation emerged from the breadth and depth-distribution of enriched metabolic markers. Nanoeukaryotes showed a profile concentrated in the epipelagic, consistent with a community whose functional specialisation is strongly anchored to the sunlit surface. Picoeukaryotes, by contrast, exhibited a greater overall number of enriched functional markers spanning nitrogen acquisition, phosphorus scavenging, and sulfur assimilation pathways across depth zones. This wider functional repertoire is consistent with the greater diversity of metabolic responses to environmental gradients documented in marine picoeukaryotes (45), and with the ecological advantages conferred by high surface-area-to-volume ratios that facilitate efficient nutrient acquisition under oligotrophic conditions (46).

Despite this size-fraction divergence, heterotrophic lineages made an increasing contribution to phosphorus and sulfur acquisition with depth in both nanoeukaryotes and picoeukaryotes, with the most pronounced signal in sulfur assimilation, where heterotrophs accounted for the large majority of bathypelagic markers. Higher absolute read counts in the pico than the nano fraction rule out size-fraction DNA leakage as the source, supporting their presence in the picoplankton size range. Carbon and nitrogen cycling markers, by contrast, remained dominated by photosynthetic and mixotrophic taxa across all depth zones, identifying P and S acquisition as the primary biogeochemical niches where heterotrophic eukaryotes emerge as significant players in the dark ocean. The persistence of these heterotrophic signals in the metatranscriptomes (Fig. S8, Table S17), together with their retention under stringent sequence-identity filtering, further supports their attribution to resident deep communities rather than sinking-derived DNA. By contrast, photosystem and Calvin-cycle transcripts likely reflect recently exported surface cells and do not provide evidence of in situ activity by resident deep-sea communities.

Phosphorus metabolism showed marked size-dependent differentiation. The surface enrichment of *PHO84* in nanoeukaryotes is consistent with well-characterised high-affinity inorganic phosphate acquisition under oligotrophic conditions (33, 47, 48). In picoeukaryotes, phosphorus scavenging markers were absent in the epipelagic but emerged with depth, shifting in the bathypelagic toward a community of carriers including Fornicata, Choanoflagellata, Foraminifera, and Ascomycota alongside residual Bacillariophyta. The restriction of Foraminifera carriers to the pico fraction is consistent with current models of foraminiferal reproduction, whereby adults sink through the water column and aggregate at depth for gamete production and release (49), suggesting that the signal may reflect this reproductive life stage. Detection of phoD transcripts assigned to Foraminifera, alongside evidence that adults accumulate phosphorus across depth zones (50), further supports a role for this group in phosphorus dynamics across life stages.

Nitrogen acquisition markers in the pico fraction spanned multiple assimilatory routes across depth zones, indicating broad N-source flexibility in the deep picoeukaryotic community. In the nano fraction, transport markers at depth were dominated by Bacillariophyta, potentially reflecting passive flux of sinking diatom biomass rather than active uptake.

Sulfur assimilation markers were enriched in the epipelagic and mesopelagic of the pico fraction, dominated by Acantharea (whose skeletons are uniquely composed of strontium sulfate, SrSO4). In the nano fraction, sulfur assimilation was restricted to the bathypelagic, with SiR enrichment driven predominantly by Choanoflagellata (Fig. S7). The sulfate assimilation pathway is conserved across aplastidic eukaryotes and operates cytosolically (51); its enrichment in predominantly heterotrophic taxa at depth points to assimilatory sulfate reduction as an underappreciated deep-ocean metabolic strategy.

N, P, and S marker enrichment at depth paralleled patterns in prokaryotic communities from the same samples (52), including a gene-level parallel in bathypelagic alkaline phosphatase (*phoD*). This convergence across phylogenetically distant domains suggests the deep ocean selects strongly for resource-acquisition flexibility regardless of domain affiliation (9).

Environmental forcing of functional composition: convergence, divergence, and niche specialisation Across both fractions, photosynthesis showed concordant negative correlations with depth and strong responses to temperature, identifying these as the principal axes of shared functional environmental response. Genetic information processing pathways were predominantly surface-associated, tracking temperature and chlorophyll positively and declining with depth.

Against this shared background, the TCA cycle and aminoacyl-tRNA biosynthesis decoupled between fractions. In the nano fraction, both correlated negatively with temperature and salinity and positively with depth and inorganic nutrients. This result is at odds with the surface enrichment of heterotrophic markers in the same fraction, and apparently contradicts the GSEA classification of these pathways as depth-invariant. The contradiction resolves when the analyses are seen as capturing different aspects of the same pattern: depth-based analyses reflect abundance changes across layers, while correlation-based GSEA captures associations arising when sinking biomass accumulates in cold, nutrient-rich waters. This points to passive export rather than depth-specific activity, aligning with the role of nano-sized plankton in vertical fluxes (53, 54). In the pico fraction, both TCA cycle and aminoacyl-tRNA pathways correlated positively with temperature and chlorophyll and negatively with depth, suggesting resident communities coupled to photic-zone production.

Each fraction also showed exclusive associations pointing to distinct niches. In the pico fraction, amino acid degradation (valine, leucine, isoleucine), butanoate, and benzoate metabolism were positively associated with depth and inorganic nutrients and negatively with temperature and chlorophyll, consistent with communities processing diverse organic compounds in the aphotic zone. Co-enrichment of secretion system and two-component signalling pathways at depth may reflect adaptations for organic matter sensing under low-carbon conditions (55). This catabolic orientation aligns with picoeukaryote genome streamlining, which preferentially eliminates biosynthetic pathways for soluble metabolites (notably amino acids) when products can be scavenged from the medium, suggesting deep pico functional profiles reflect both ecological adaptation and an evolutionary legacy of genome reduction (56).

In the nano fraction, pyruvate, sulfur, and glyoxylate/dicarboxylate metabolism were positively associated with depth and inorganic nutrients. The glyoxylate cycle synthesises carbohydrates from fatty-acid-derived acetyl-CoA (57, 58), conserving carbon when labile carbohydrates are scarce. As sinking POM is progressively stripped of labile carbohydrates and proteins, leaving lipid-enriched residues at depth (59), this pathway provides a route to recover carbohydrate carbon. Restriction to the nano fraction is consistent with greater exposure to lipid-enriched sinking POM in this size class (53, 60), though the 3–20 µm fraction also captures particle-attached smaller cells, a distinction unresolvable from size-fractionated metagenomes.

### Projected responses of eukaryotic metabolic potential to ocean warming

Under SSP5–8.5 (mean projected warming: ∼2.2 °C epipelagic, ∼1.0 °C mesopelagic), GAM projections of picoeukaryote functional potential revealed spatially structured responses to end-of-century warming, with epipelagic and mesopelagic shifts and a stable bathypelagic. Projections were restricted to the pico fraction, as temperature-only GAMs for the nano fraction yielded low explanatory power, potentially because the passive export signal reflects downward transport more strongly than environmental conditions.

The dominant response, an increase in photosynthesis-related potential in subtropical epipelagic waters, with additional increases in the mesopelagic western North Atlantic, is consistent with these communities’ thermal performance position, where warming stimulates Calvin cycle and electron transport rates (61), and with predicted expansion of photosynthetic picoplankton in warming subtropical gyres (27). The mesopelagic signal likely reflects downward export rather than photosynthesis at depth, as deep chlorophyll maxima remain confined to the lower euphotic zone even under increased stratification (62). Porphyrin metabolism showed similar patterns, pointing to coordinated increases in photosynthetic and respiratory electron transfer capacity.

The projected increases in TCA cycle and aminoacyl-tRNA biosynthesis point to enhanced aerobic carbon metabolism and biosynthetic capacity in cold, productive waters. Their concentration in subpolar surface waters and the North Atlantic mesopelagic is consistent with predicted temperature-driven increases in high-latitude phytoplankton production(26, 27), and with the positive associations of these pathways with productivity indicators (Fig. 4). Together, these patterns suggest that warming may enhance both carbon processing and the translational machinery required to support increased biomass production in productive high-latitude regions.

The most striking responses were declines in transport functions and heterotrophy markers, spatially concentrated where photosynthetic and aerobic potential increases. Transport markers decline in the epipelagic and mesopelagic, suggesting warming-driven compositional shifts reduce demand for active nutrient scavenging as phototrophy dominates. Heterotrophy markers showed the largest declines. Taxonomic decomposition revealed that these markers are contributed by a mixed community spanning all major trophic categories (Fig. S12): strict phototrophs, mixotroph-capable lineages, and obligate heterotrophs, each contribute a substantial share. The β-oxidation markers’ negative correlation with temperature is consistent with shifts toward heterotrophic transcriptional profiles at greater depths in protistan metatranscriptomes (37, 63, 64). Notably, mixotroph-capable lineages contribute substantially to both the heterotrophy and increasing photosynthesis signals (Fig. S12B), which show opposing spatial patterns (Fig. 5). Together, these observations are consistent with within-lineage metabolic rebalancing in mixotrophic taxa contributing to the projected decline in heterotrophy, alongside compositional shifts among strict phototrophs and obligate heterotrophs. Because pathway abundance represents the aggregate contribution of many taxa rather than the physiology of any single organism, compositional turnover and temperature-driven shifts in the distribution of thermal traits among contributing taxa can smooth or obscure the decline phase expected for individual thermal performance curves (65).

The consistency of projected functional shifts across independent environmental predictors and their robustness to model-flexibility choices lend support to these interpretations and argue against overfitting as an explanation (Table S16). Single-predictor GAMs for NO₃ and Si agreed in direction with the temperature-driven projections wherever all three predictors reached significance simultaneously (Fig. S11), although this concordance was less stable at depth, where significant-pixel counts were lower. Temperature- and nutrient-driven projections were nonetheless only weakly to moderately spatially correlated, even within a single climate ensemble, indicating that while the direction of projected functional change is robust to predictor choice, its precise geographic expression depends in part on which aspect of climate change dominates locally.

Collectively, these responses suggest that warming will narrow the metabolic repertoire of picoeukaryote communities toward phototrophic dominance, with consequences for trophic transfer efficiency, the composition of exported organic matter, and the biological carbon pump.

### Methodological considerations and limitations

The ∼79% reduction in predicted eukaryotic ORFs following curation highlights the substantial risk of prokaryotic signal inflation in eukaryotic metagenomic analyses of small size fractions, where prokaryotes dominate sequencing reads (66). Without stringent filtering, prokaryotic sequences misclassified as eukaryotic would disproportionately inflate functions shared across domains while leaving eukaryote-specific pathways relatively unaffected, systematically biasing the functional profile toward a prokaryote-like signature. The magnitude of this reduction underscores the importance of multi-step curation in eukaryotic metagenomic datasets and suggests that functional profiles derived from small size-fraction metagenomes without comparable filtering should be interpreted with caution (67). Additionally, the curation pipeline retained only ORFs matching eukaryotic references across all three databases (EukProt, MarFERReT, and MMETSP). As MMETSP is predominantly composed of cultured photosynthetic organisms (68), this intersection filter may systematically underrepresent ORFs from heterotrophic taxa with limited reference sequences. The heterotrophic metabolic strategies documented in this study, including the contributions of Choanoflagellata, Foraminifera, Acantharea, and fungi to deep-ocean nutrient cycling, should therefore be considered conservative estimates that emerged despite this bias. The direction of this bias is supported by comparison with 18S rDNA mTags recovered from the same metagenomes (Table S14, Fig. S1). Phototrophs dominated the ORF-based catalogue but represented only a minor fraction of mTag reads across all depths, whereas amplicon-defined environmental clades such as MALV (Syndiniales) and RAD (radiolaria), which lack protein-level references, accounted for 17–23% of epipelagic mTag reads yet were absent from our catalogue. Other heterotrophic lineages are present in our catalogue but coarsely resolved: NCBI groups Labyrinthulomycetes with Bicosoecida and other Opalozoa under Bigyra, which could only be mapped to Bicosoecida; most MAST lineages lack a ranked NCBI taxonomy and only MAST-3 and MAST-4 could be recovered; and Syndiniales are nested within Dinophyceae. These contrasting biases reflect fundamental differences between the approaches: ribosomal surveys over-represent high rDNA copy-number taxa such as radiolaria and dinoflagellates, whose marker abundance scales with cell and genome size (69), whereas ORF-based catalogues are constrained by reference coverage but better recover diatoms. The two approaches thus offer complementary but biased views that together bound the likely range of true community composition (33, 67); and while our catalogue composition is influenced by reference bias, the recovery of Foraminifera, Choanoflagellata, and Cercozoa shows higher representation than in mTags, indicating that the deep-ocean heterotrophic signal is not a methodological artefact.

The GAM-based climate projections were fitted to vertical profile data from tropical and subtropical stations and were subsequently extrapolated to a global geographic context. The resulting spatial patterns should therefore be interpreted as indicative of the direction and relative magnitude of potential responses rather than precise regional forecasts. Extending this framework to higher-latitude datasets, where thermal structure differs substantially from the training data, will be necessary to improve projection robustness. The absence of significant bathypelagic projections likely reflects the limited magnitude of projected temperature change at depth within the CMIP6 ensemble by 2100, which remained within the thermal range sampled during the Malaspina expedition and thus within the domain of GAM predictive reliability. Because temperature, nutrients, and depth are strongly intercorrelated, temperature-only projections cannot distinguish a thermal effect from the broader environmental gradient with which temperature co-varies (Methods; Table S9). Compounding this, the temperature–nutrient collinearity structure itself weakens moderately in the epipelagic and substantially at depth between the training and projected periods (Table S10), consistent with warming-driven stratification decoupling surface temperature from nutrient supply. Projected changes should therefore be read as tracking this broader gradient rather than isolating a temperature-specific mechanism, particularly outside the epipelagic. Significant projected pixels for all predictors fell within the well-sampled core of the training data (Table S8). We note that SSP5-8.5 is no longer considered a plausible high-end scenario within the CMIP7 framework (70). Here, we use it to maximise signal detectability; projected magnitudes should therefore be interpreted as upper bounds, while the direction and spatial pattern of the response are more scenario-robust.

Our curated catalogue and size-fraction-resolved framework establish a resource for future investigations of eukaryotic functional ecology in the global ocean. We provide a functionally resolved view of pico- and nano-sized microbial eukaryote communities in the subtropical ocean across the entire water column, revealing that depth together with cell size structure eukaryotic functional potential. Fraction-specific enrichment patterns, environmental response profiles, and the contrasting explanatory power of environmental models across size fractions point to a fundamental ecological distinction: picoplankton communities harbour more depth-adapted resident assemblages, while nanoplankton functional composition at depth is more influenced by passive export of surface-derived biomass (Fig. 6). Heterotrophic protists, including Choanoflagellata, Foraminifera, and Acantharea, emerge as undercharacterised contributors to phosphorus and sulfur cycling in the dark ocean. Bathypelagic metatranscriptomes show that this potential is expressed: heterotrophic lineages produced most transcripts of phosphate transporters and sulfur-assimilation genes, which remained detectable under stringent mapping, consistent with metabolically active resident communities. Functional projections under end-of-century warming suggest a narrowing of metabolic repertoire toward phototrophic dominance, consistent with metabolic rebalancing within mixotrophic taxa, with potential consequences for trophic transfer efficiency and the composition of exported organic matter.

**Fig. 6.**
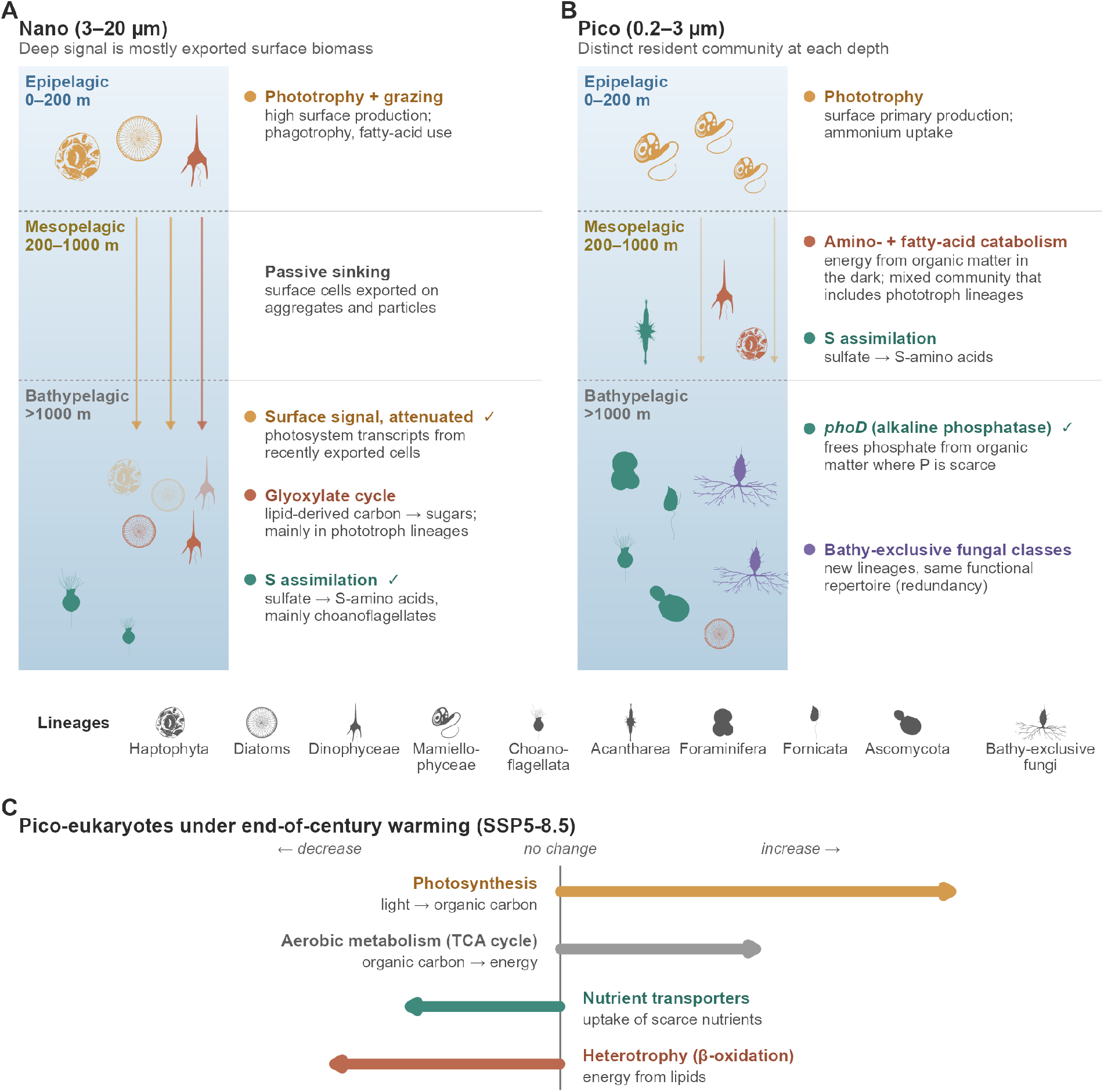
Conceptual summary of depth-related functional patterns of marine microbial eukaryotes and their projected response to warming. Positions and sizes are not quantitative. Silhouettes represent lineages discussed in the text, coloured by the function they illustrate (Yellow: phototrophy; Red: heterotrophic metabolism; Green: P and S acquisition; Purple: fungal lineages). ✓: function also detected in bathypelagic metatranscriptomes. (A) Nano fraction (3– 20 µm): surface lineages and their export to depth (arrows; faded silhouettes). (B) Pico fraction (0.2–3 µm): lineages and functions characteristic of each depth zone; faint arrows indicate minor export. (C) Projected direction of change in pico-fraction metabolic potential in epi- and mesopelagic waters under SSP5-8.5 by the end of the century; arrow lengths are schematic.

## MATERIALS AND METHODS

### Sample collection

A total of 116 water samples were taken during the Malaspina 2010 expedition (http://www.expedicionmalaspina.es) on board the R/V Hespérides from January 4th to July 5th, 2011, corresponding to 11 sampling stations ( Fig. 1A-B, Table S1). Each station was profiled by collecting water from 7 discrete depths (except station 49, with 6 depths) from the surface (3 m) to the bathypelagic layer, down to 4,000 m. Water samples were collected either with a rosette of Niskin bottles (12 L each) mounted on a CTD rosette system or with a large Niskin bottle (30 L) for surface sampling. Approximately 12 L of seawater were prefiltered through a 200 μm nylon mesh and then sequentially filtered using a peristaltic pump through a 20 μm nylon mesh, followed by 3 μm and 0.2 μm pore-size 142 mm Millipore polycarbonate filters. Filters were immediately flash-frozen in liquid nitrogen and stored at –80 °C until processing in the laboratory.

### DNA Extraction

DNA extracts were obtained with the phenol-chloroform protocol (71). For the nanoeukaryotic fraction (3-20 μm) we obtained 40 samples, and for the picoeukaryotic fraction (0.2-3 μm) we obtained 76 samples (Fig. 1 A-B, Supp. Table 1). The extraction protocol was previously reported in (52). Briefly, filter membranes were first sectioned into small fragments using sterile razor blades and resuspended in 3 mL of lysis buffer (40 mM EDTA, 50 mM Tris-HCl, 0.75 M sucrose). Samples were incubated at 37 °C for 45 min in lysis buffer supplemented with lysozyme (final concentration 1 mg mL⁻¹) under gentle agitation. Subsequently, sodium dodecyl sulfate (SDS; final concentration 1%) and proteinase K (final concentration 0.2 mg mL⁻¹) were added, and incubation was continued at 55 °C for 60 min with gentle mixing.

The resulting lysate was subjected to phenol–chloroform extraction. An equal volume of Phenol:Chloroform:Isoamyl alcohol (25:24:1, v/v/v) was added, mixed thoroughly, and centrifuged for 10 min at 3,000 rpm. The aqueous phase was recovered, and the extraction step was repeated once. To remove residual phenol, the recovered aqueous phase was further extracted with an equal volume of Chloroform:Isoamyl alcohol (24:1, v/v), followed by centrifugation and collection of the aqueous phase.

DNA was then concentrated using a Centricon centrifugal concentrator (Amicon Ultra-4, Millipore, Ultracel-100 membrane). The concentrate was washed three times by adding 2 mL sterile Milli-Q water per wash to remove potential contaminants that could interfere with downstream library preparation. Finally, the DNA extract was reduced to a final volume of 100–200 µL.

### Sequencing

Whole metagenome shotgun sequencing was performed at CNAG (Barcelona, Spain; http://cnag.cat/). Short-insert paired-end libraries were prepared with the Illumina TruSeq Sample Preparation kit (Illumina Inc.) for all picoeukaryotic and nanoeukaryotic samples. Sequencing was carried out on HiSeq 2000/4000 Illumina platforms for all samples, with a read length of 2 × 101 or 2 x 150 bp (16, 52). Sequencing yielded an average of 27.0 ± 19.2 Gbp per sample (mean ± standard deviation).

### Metagenomic assembly, gene prediction, and functional annotation

A total of 116 metagenomic FASTQ files were quality-checked, cleaned, and trimmed before assembly using cutadapt v1.16 (72). Each metagenome was individually assembled using MEGAHIT v1.1.3 (73) with the --presets meta-large option, which is recommended for large and complex metagenomes. Following assembly, contigs were renamed and processed with EukRep v0.6.5 (74) using default parameters to separate eukaryotic and prokaryotic contigs. Gene prediction on the eukaryotic contigs was performed using MetaEuk v1-ea903e5, using reference gene profiles from MMETSP, uniclust50, and MERC (75). All predicted genes were concatenated into a single dataset. This resulted in 63,684,194 redundant predicted genes (enriched in eukaryotes, but including prokaryotes). To remove redundancy, clustering was performed using Linclust (76) from MMseqs2 v9-d36de at 95% sequence identity thresholds with 80% coverage of the shorter fragments. The non-redundant 95% catalogue contained 33,142,301 open reading frames (ORFs) longer than 250 bp. This minimum length was considered to avoid very short genes.

Functional annotation of the predicted ORFs was carried out using DIAMOND diamond/0.9.22 (77) and blastp (78) against the KEGG database (kegg_release_20170220) (79).

### Microbial eukaryotic gene catalogue curation

We observed that the initial gene catalogue contained prokaryotic ORFs. To ensure a reliable assessment of the metabolic potential of microbial eukaryotes in the water column, we further curated the catalogue using an ad hoc multi-step process. Firstly, we performed three parallel taxonomic assignments using the ‘easy-taxonomy’ module from MMseqs2 (mmseqs2/13-45111) (80), against three different databases: EukProt (81), MarFERReT v1.1.1 (82), and MMETSP_Uniclust90. The first two databases contain eukaryotic reference sequences, whereas the third is a custom reference database assembled from the MMETSP (The Marine Microbial Eukaryote Transcriptome Sequencing Project) (68) and Uniclust90 (83) databases and includes both prokaryotic and eukaryotic references.

Only ORFs with hits in all three databases (n=17,348,923) were retained for the downstream curation steps. This filtering was necessary to remove ORFs with hits that could not be compared and reliably assigned to eukaryotes. Next, we selected the best taxonomic hit. To this end, we developed a metric termed the Tax ID score, calculated as a weighted sum of query coverage and percent identity (pident). Query coverage (defined as alignment length/query length) was assigned a weight of 0.6, while pident was assigned a weight of 0.4. Query coverage was weighted more heavily than pident because a hit spanning a larger proportion of the query sequence provides a more reliable basis for taxonomic assignment than a highly similar but shorter alignment, which may reflect conserved domains shared across distantly related taxa. For each ORF, only eukaryotic hits (Domain=Eukaryotes) with an e-value < 1e–5 and the highest Tax ID score were retained. The results comprised 8,139,291 putative eukaryotic ORFs, which were further validated against the Marine Atlas of Tara Oceans Unigenes (MATOU) reference protein database (https://www.genoscope.cns.fr/tara/), derived from eukaryotic metatranscriptomics data (22) using the MMseqs2 search tool. Only hits with e-value < 1e-5 were retained. This resulted in a total of 7,714,686 eukaryotic ORFs retained.

Next, the taxonomic assignment of the curated eukaryotic ORFs was refined by deriving a minimum consensus taxonomy for each gene that passed the filtering process and shared the same Tax ID score across multiple hits. To achieve this, we retrieved the complete taxonomic lineage (NCBI Taxonomy IDs) for each hit using the taxize R package v 0.10.0 (84), specifying the ‘ncbi’ taxonomic identifier option. The taxonomic paths were then compared and truncated to the lowest common taxonomic level (consensus taxon) shared among the hits. If no consensus was found, only the domain-level assignment was retained.

As a final step, we removed from the catalogue eukaryotic ORFs assigned to taxa outside the scope of our investigation (e.g., metazoa, plants, and macroalgae). A full list of the excluded taxa is provided as supplementary material (Table S2).

### Abundance tables curation

ORF count tables for the pico and nano size fractions were normalized in two steps. First, counts were divided by ORF length (in kilobases) to correct for gene length variation within samples. Then, values were normalized by total metagenome size to account for differences in sequencing depth among samples. This two-tier normalization approach ensured accurate and comparable gene abundance estimates across samples. The resulting normalized ORF tables were then used to generate KEGG-annotated count tables by adding the abundances of individual ORFs annotated to different KOs.

### Environmental data

Temperature and salinity vertical profiles from CTD in-situ measurements were integrated with nutrient (PO₄, Si, NO₃) and oxygen profiles obtained from the biogeochemical hindcast of the Global Ocean Mercator product, which provides 3D biogeochemical fields from 1993 onward at 1/4° horizontal resolution and 75 vertical levels (85) (Table S3), chosen over in-situ measurements for their near-complete spatial coverage across the sample set (98% vs. 80–94% for in-situ).

### Statistics

We calculated accumulation curves using a permutation-based presence–absence approach implemented in a custom R script (86). The relationship between taxonomic diversity (NCBI TAXID) and functional diversity (KOs and ORFs) was assessed using Kendall’s correlation, as implemented in the ggpubr R package v 0.6.2 (87), and visualized with the ggscatter function from the same package. Depth-specific functional and taxonomic distinctiveness were determined by identifying ORF and KO classes unique to each depth zone within each station.

To test for depth-zone differences in KO abundance, KO count tables were converted to relative abundances and centered-log-ratio (CLR)-transformed after addition of a 1×10⁻⁶ pseudocount. For each KO, a Kruskal–Wallis test was applied to CLR values across depth layers; p-values were corrected across all KOs by the Benjamini–Hochberg procedure and KOs with adjusted p < 0.05 were considered differentially abundant and further analyzed. For these, Dunn’s post-hoc test with Bonferroni correction identified pairwise differences between depth zones using dunn.test() from the R package dunn.test v1.3.6. The direction of enrichment was inferred by comparing median relative abundances across groups. An analogue procedure was applied to prokaryotic counts from the same samples previously published by (52). Finally, among the KOs showing significant variation, we identified those that could serve as metabolic markers to characterize shifts in ecological strategies across domains, size fractions, and depth zones. The list of microbial eukaryote metabolic markers was compiled from (33, 88, 89), while the prokaryotic markers were taken from (90) (Table S4).

In parallel, we identified metabolic pathways that remained constant in terms of abundance along the water column. Using hellinger-transformed count matrices, we calculated the coefficient of variation (CV) across depth zones for each individual KO. Next, we ranked KOs based on the z-scores of the coefficient of variation and performed a gene set enrichment analysis (GSEA) using the GSEA function in the clusterProfiler v 4.12.6 R package (91) with KEGG BRITE functional categories as predefined gene sets. The resulting normalized enrichment scores (NES) indicated both the direction and strength of association between BRITE pathways and CV, with higher positive values reflecting enrichment of pathways that varied strongly across depth zones, and negative NES values indicating metabolic pathways that were more constant across depth zones.

To investigate the relationship between microbial eukaryotic metabolic potential and environmental variables, we computed Kendall’s correlations between each KO (normalized to relative abundance and log₂-transformed) and the measured environmental parameters using the cor.test function in R. KOs with significant correlations (p < 0.01) were retained. For each environmental variable, we then performed gene set enrichment analysis on a ranked list of KOs ordered by their Kendall’s τ values, using the GSEA function in the clusterProfiler v 4.12.6 R package (91) and KEGG BRITE functional categories as predefined gene sets. The resulting normalized enrichment scores (NES) indicated both the direction and strength of association between BRITE pathways and environmental gradients, with positive NES values reflecting pathway enrichment at higher environmental values and negative NES values indicating enrichment at lower values. Finally, we selected the top 30 enriched pathways based on the absolute magnitude of their NES values.

### ORF–mTag composition comparison

To assess how reference-database composition shaped the catalogue, we compared ORF-based taxonomy (NCBI) with 18S rDNA mTags from the same metagenomes, extracted and classified against PR2 following Obiol et al. (2020). NCBI class-level assignments were mapped to PR2 groups with a manually curated crosswalk (Table S5), and per-group relative abundances were compared across depth zones (epipelagic ≤200 m, mesopelagic 200–1000 m, bathypelagic >1000 m). Because Syndiniales (PR2 MALV-II) are nested within Dinophyceae in NCBI and cannot be separated on the ORF side, MALV-II was collapsed into Dinoflagellata on both sides; ORF Dinoflagellata also include Syndiniales group IV (Hematodinium), which remains separate (MALV-IV) in mTags. MALV-I/III/V and RAD clades have no NCBI protein references and appear only in mTags. Notably, ORF and mTag abundances reflect different currencies (i.e., gene content versus rDNA copy reads) and were compared as compositional profiles, not estimates of a common quantity.

### Metatranscriptomic validation of depth-enriched markers

To test whether the eukaryotic metabolic markers found differentially abundant in the bathypelagic metagenomes reflect active metabolism rather than passive accumulation of sinking material, we examined their expression in an independent set of 51 bathypelagic metatranscriptomes from the same expedition (Fig. 1; Table S6) (92). Marker expression was quantified using two complementary approaches against the KO marker ORF sequences (Table S4). First, transcript abundances were estimated with Salmon v1.5.2 (93) in selective-alignment mode (--validateMappings), with automatic library-type detection (-l A) and a k = 31 index and expressed as transcripts per million (TPM); markers with TPM > 0 were considered transcribed. Second, to confirm that these signals were not artefacts of permissive mapping or cross-mapping from prokaryotic transcripts, reads were aligned with BWA-MEM2 v2.2.1 (94), retaining only primary alignments with mapping quality ≥ 10, and per-marker coverage was computed with CoverM v0.7.0 (95) under stringent filtering: ≥95% read identity, ≥80% of each read aligned, and a minimum covered fraction of 75% of the marker length. Persistence of marker detection under these criteria, where spurious and cross-domain mapping is minimized, was taken to indicate genuine eukaryotic marker transcription in the bathypelagic.

### Temperature–function modelling

To quantify the relationship between temperature and microbial eukaryotic functional potential, we modeled the association between temperature and selected metabolic pathways using generalized additive models (GAMs). Specifically, we examined the relationship between in situ sea temperature and the log-transformed relative abundance of KEGG orthologs (KOs) aggregated from those showing significant normalized enrichment scores in the temperature-based GSEA analysis across both size fractions. We restricted the analysis to KOs belonging to a predefined set of metabolic pathways representing ecologically relevant functions, including photosynthesis, porphyrin metabolism, the citrate cycle (TCA), aminoacyl-tRNA biosynthesis, and transporters, as well as selected BRITE pathways and heterotrophy-related KO markers. Analyses were conducted separately for the two size fractions: pico- (0.2–3 µm) and nano-plankton (3–20 µm). Read counts were converted to relative abundance. Values were then log₂(x + 1) transformed, and ortholog-level abundances were aggregated at the BRITE category level for each sample. For each BRITE category and size fraction, a separate GAM was fitted using the mgcv v1.9.1 package in R (96). Functional abundance was modelled as a smooth function of temperature:

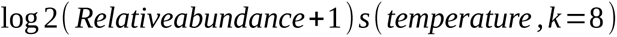

Models were estimated using restricted maximum likelihood (REML). Model performance was assessed using adjusted R^2^ and effective degrees of freedom (EDF), and model assumptions were evaluated using diagnostic checks implemented in gam.check(). Derivative confidence intervals were calculated using the R package gratia v 0.11.2( Simpson, 2024) and were used to identify, for each fitted smooth, the sub-ranges of the training data over which the slope was statistically distinguishable from zero, and thereby to test whether apparent non-monotonicity in point estimates reflected significant reversals rather than noise. As a secondary check on sensitivity to model flexibility, each model was also refitted at a reduced basis dimension (k=5) and the resulting shape and effective degrees of freedom compared against the primary k=8 fit (Table S7). Because GAMs fitted to the nano-plankton size fraction showed consistently low explanatory power (adjusted R^2^), all downstream spatial projections and interpretations were restricted to pico-plankton models. Monthly temperature projections for the epipelagic, mesopelagic, and bathypelagic layers under the SSP5–8.5 scenario were obtained from (97). Raster time series were reduced to climatological means by computing annual averages and aggregating across two periods: 2015–2020 (baseline) and 2095–2100 (future). Each GAM was projected onto the processed rasters to generate spatial predictions of functional potential under present and future conditions.

To prevent extrapolation, predictions were limited to the empirical temperature range of each model. The proportion of significant projected pixels falling within the dense (5th–95th percentile) core of the training predictor distribution, as opposed to its sparser tails, was quantified for each predictor and depth zone (Table S8).

Prediction uncertainty was propagated by combining baseline and future standard errors:

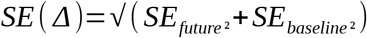

To aid interpretation, significant log-scale differences were then back-transformed to the natural scale and expressed as percentage change using:

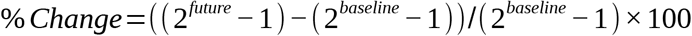

Final maps, therefore, represent only robust, temperature-associated shifts in microbial functional potential across ocean depths.

### Corroboration with nutrient predictors

To assess whether the projected functional shifts were specific to temperature forcing or reflect a broader response along the oligotrophic–eutrophic gradient, we fitted independent single-predictor GAMs for nitrate (NO₃) and silicate (Si) on the same pico-plankton training data, using the model structure described above. Prior to fitting, predictor collinearity was assessed using pairwise Pearson correlations. Temperature, NO₃, and Si were strongly intercorrelated (temp×NO₃ r = −0.89, temp×Si r = −0.72, NO₃×Si r = 0.81), confirming their shared role as proxies of the same warm/oligotrophic to cold/eutrophic oceanographic gradient (Table S9). To evaluate whether this collinearity structure was preserved under the projected scenario, the same pairwise correlations were recomputed across the projected (2095–2100) CMIP6 fields for each depth zone and compared against the training-period values (Table S10). Phosphate (PO₄) was excluded as redundant with NO₃ (NO₃×PO₄ r = 0.98). A combined multi-predictor GAM was not fitted, as the high intercorrelation among predictors induces concurvity which inflates standard errors and renders smooth estimates unreliable (Table S11). Each nutrient predictor was therefore modelled separately, as an independent lens on the same underlying biological signal (Table S12).

Future temperature, NO₃ and Si forcing were obtained from the same CMIP6 ensemble of seven Earth system models under SSP5–8.5 (CESM2-WACCM, CMCC-ESM2, CNRM-ESM2-1, IPSL-CM6A-LR, MPI-ESM1-2-HR, MPI-ESM1-2-LR, UKESM1-0-LL; Table S13). For each model, native vertical levels were assigned to epipelagic (0–200 m), mesopelagic (200–1,000 m), and bathyabyssopelagic (>1,000 m) depth zones (Table S13) and output was regridded to a common 0.5° spatial grid. Within each depth zone, values were averaged across vertical levels, and an ensemble mean was then computed across available models for each variable, depth zone, and time period (baseline: 2015–2020; future: 2095–2100). Nutrient GAM projections were generated following the same pipeline as the temperature-based projections described above; ensemble nutrient concentrations were rescaled from mol m^⁻3^ to mmol m^⁻3^ to match the units of the station training data prior to projection.

Agreement between predictors was assessed in two complementary ways. First, spatial coherence was quantified as the Pearson correlation between projected percentage-change surfaces, computed both before and after significance masking, because masked correlation becomes unreliable where significant pixels are spatially sparse. Second, directional concordance was quantified as the percentage of pixels (among those where at least one predictor showed significant projected change) at which all three predictors yielded significant change of the same sign. Sensitivity of directional concordance to model flexibility was assessed by recalculating concordance using the k=5 fits described above (Table S16). Directional concordance, rather than masked spatial correlation, was used as the primary criterion for agreement between temperature and nutrient forcing.

## Acknowledgments

We thank the R/V Hespérides captain and crew, the chief scientists in the Malaspina expedition legs, and all project participants. Bioinformatics analyses were performed at the MARBITS platform of the Institut de Ciències del Mar (ICM) [https://marbits.icm.csic.es] and Centro de Supercomputación de Galicia (CESGA; https://www.cesga.es/).

## Author contributions

Conceptualization: R.L., M.G.

Data Curation: M.G., L.M., F.L., C.R.G.

Methodology: R.L, M.G., R.B.

Investigation: S.G.A., S.A., C. M. D., R. M., C. R.G., M.G., R.L.

Methodology: M.G., R.L. Formal

Analysis: M.G., R.L.

Visualization: M.G.

Supervision: R.L.

Writing—original draft: M.G.

Writing—review & editing: R.L., R.M., C.M.D., R.B., F. L.

## Data, code, and materials availability

Raw metagenomic and metatranscriptomic reads from the Malaspina 2010 expedition are available at ENA under accession number PRJEB46336. The eukaryotic gene catalogue, together with the ORF and KO count matrices will be available on Zenodo. Supplemetary tables S1-S17 are available on Zenodo (https://doi.org/10.5281/zenodo.23567862).

## Supplementary Figures

**Fig. S1.**
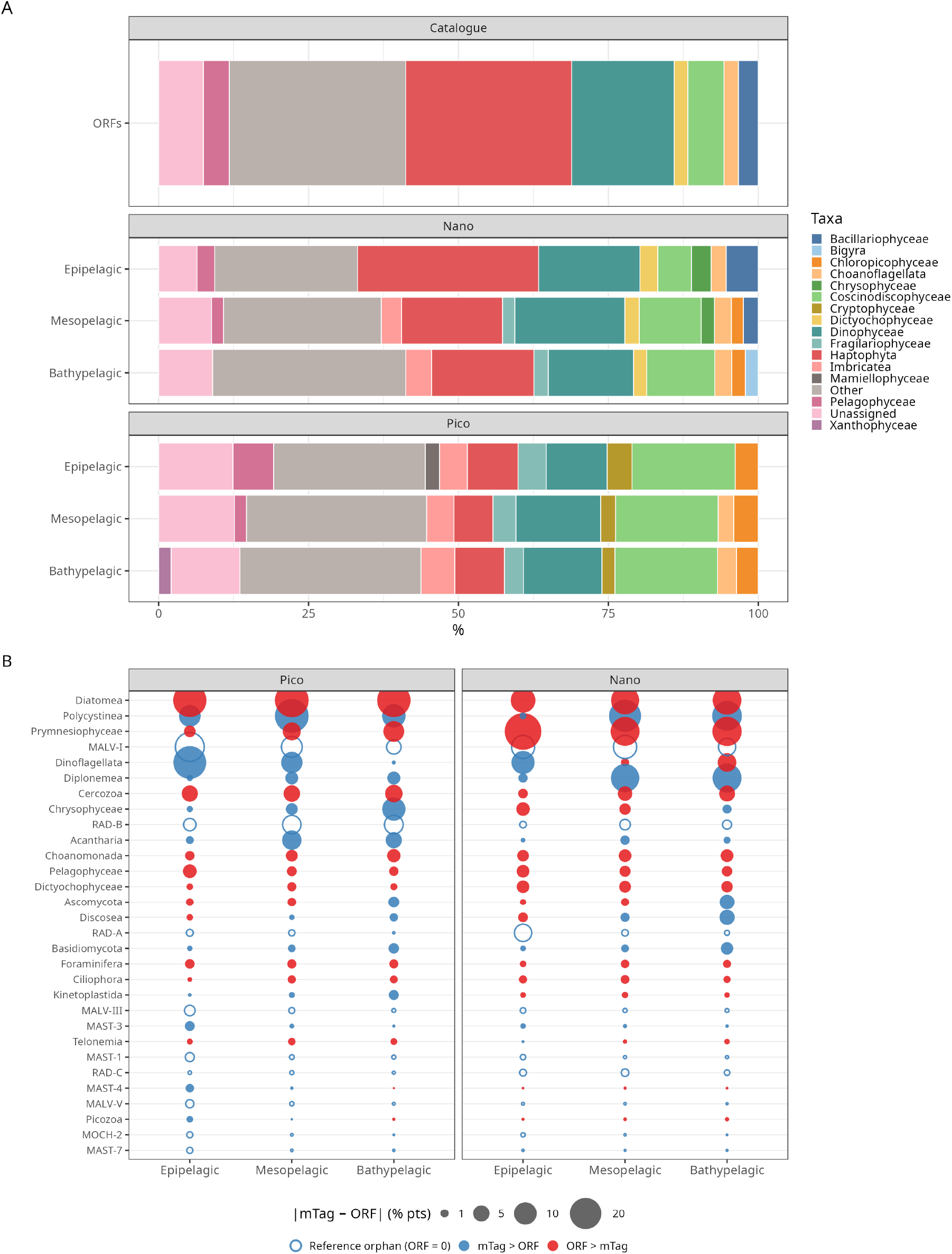
Taxonomic composition of the ORF catalogue and comparison with 18S rDNA mTags. (A) Taxonomic composition of the ORF catalogue as a percentage of total ORFs (top row), and mean relative abundance of taxonomic groups per depth zone and size fraction. (B) Difference between ORF and mTag mean relative abundance per PR2 group, depth zone and size fraction (pico, 0.2–3 µm; nano, 3–20 µm). Bubble area is proportional to |mTag − ORF| (%); groups are ordered by mean discordance across depth zones and size fractions. Hollow circles: groups with no ORFs in the catalogue (ORF = 0); filled blue: mTag > ORF; filled red: ORF > mTag. Taxonomic mapping and per-group values are given in Tables S5 and S14; see Methods for details.

**Fig. S2.**
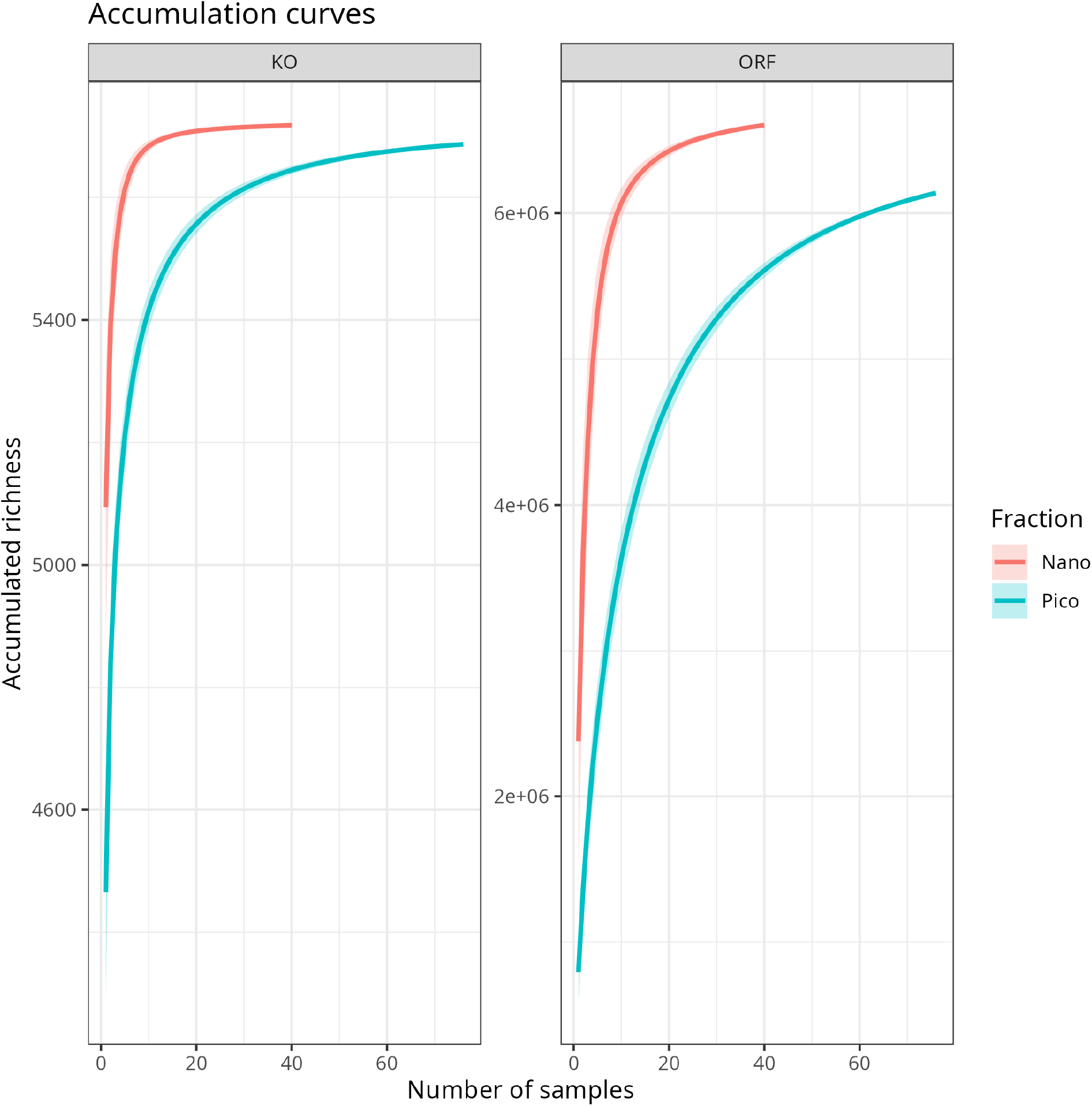
Sample-based accumulation curves for ORF and KO richness in pico (0.2–3 µm) and nano (3–20 µm) size fractions. Accumulated richness of unique ORFs (right panel) and KEGG Orthologs (KOs; left panel) as a function of the number of samples added, for the pico (n = 76) and nano (n = 40) size fractions. Curves represent the mean accumulated richness across 100 random permutations of sample order; shaded ribbons indicate ±1 standard deviation.

**Fig. S3.**
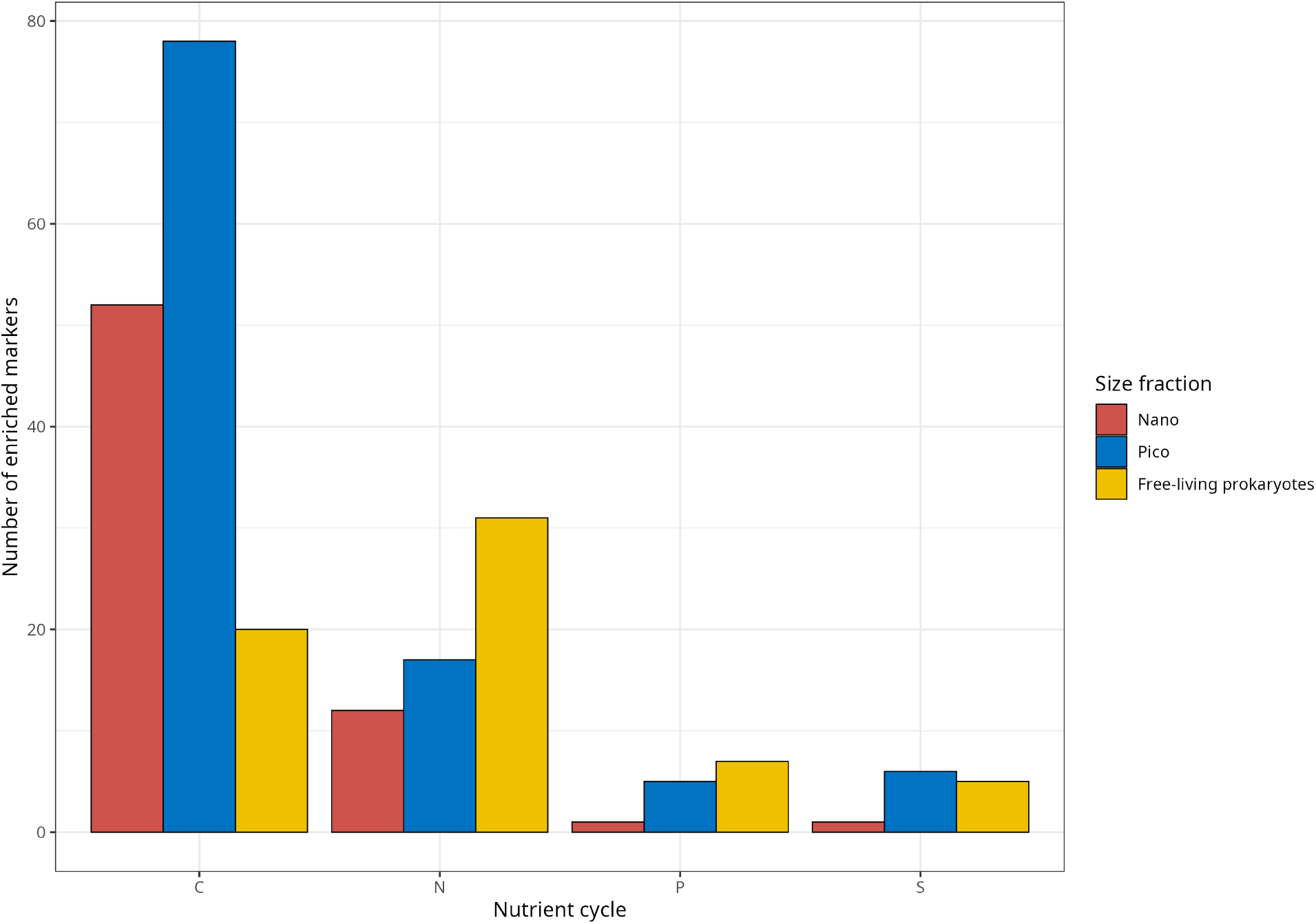
Total number of significantly depth-enriched biogeochemical markers per nutrient cycle and size fraction. Total count of unique KOs significantly enriched across depth zones (Kruskal-Wallis, Dunn post-hoc p < 0.01) for each nutrient cycle (C, carbon; N, nitrogen; P, phosphorus; S, sulfur) in nano- and pico-eukaryotic size fractions and free-living prokaryotes. Counts reflect all enriched markers regardless of the direction or depth zone of enrichment.

**Fig. S4.**
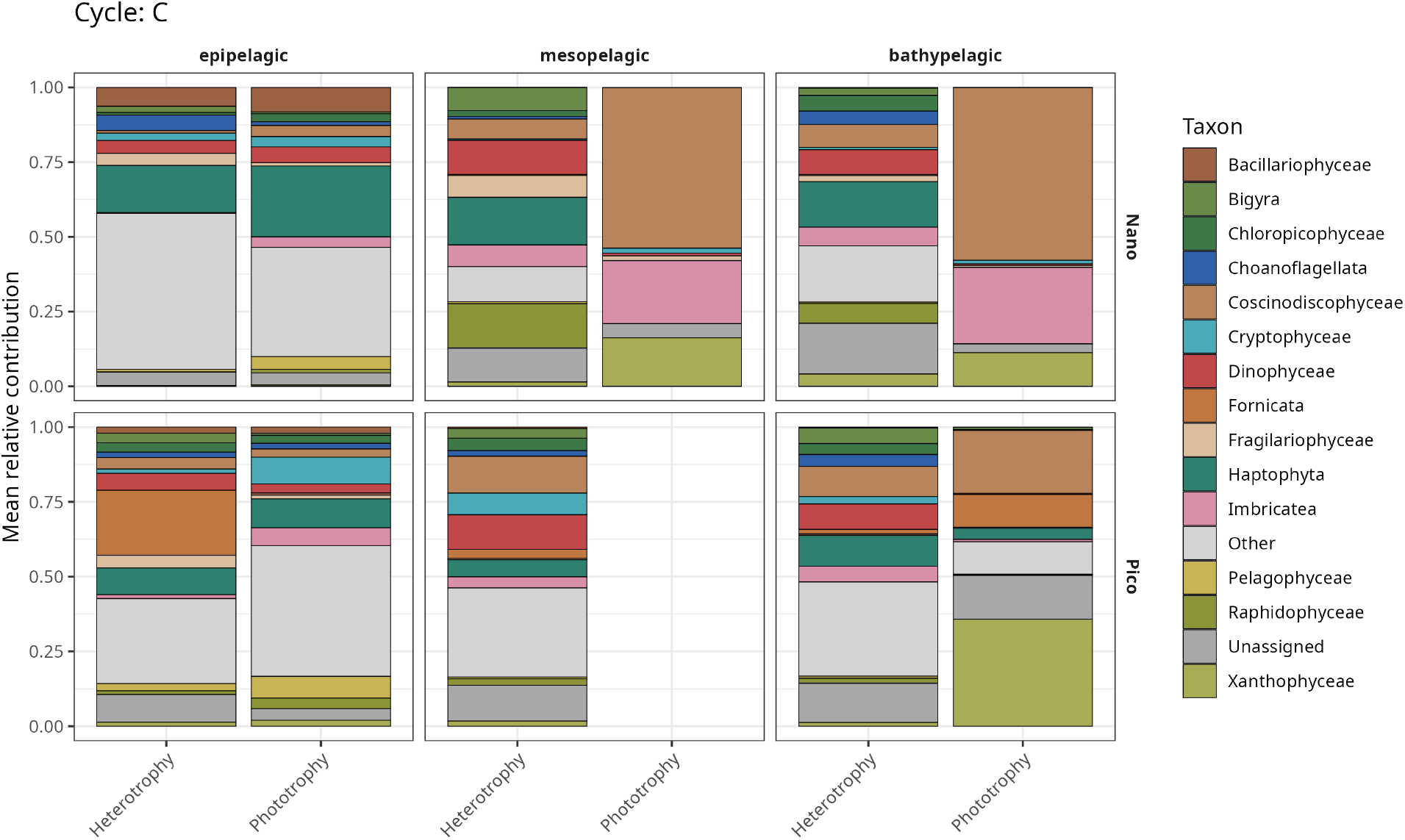
Taxonomic contribution to enriched carbon cycling markers. Stacked barplots showing the mean relative contribution of the top eukaryotic taxa to significantly depth-enriched carbon (C) cycling markers, for nano- and pico-size fractions across epipelagic, mesopelagic, and bathypelagic depth zones

**Fig. S5.**
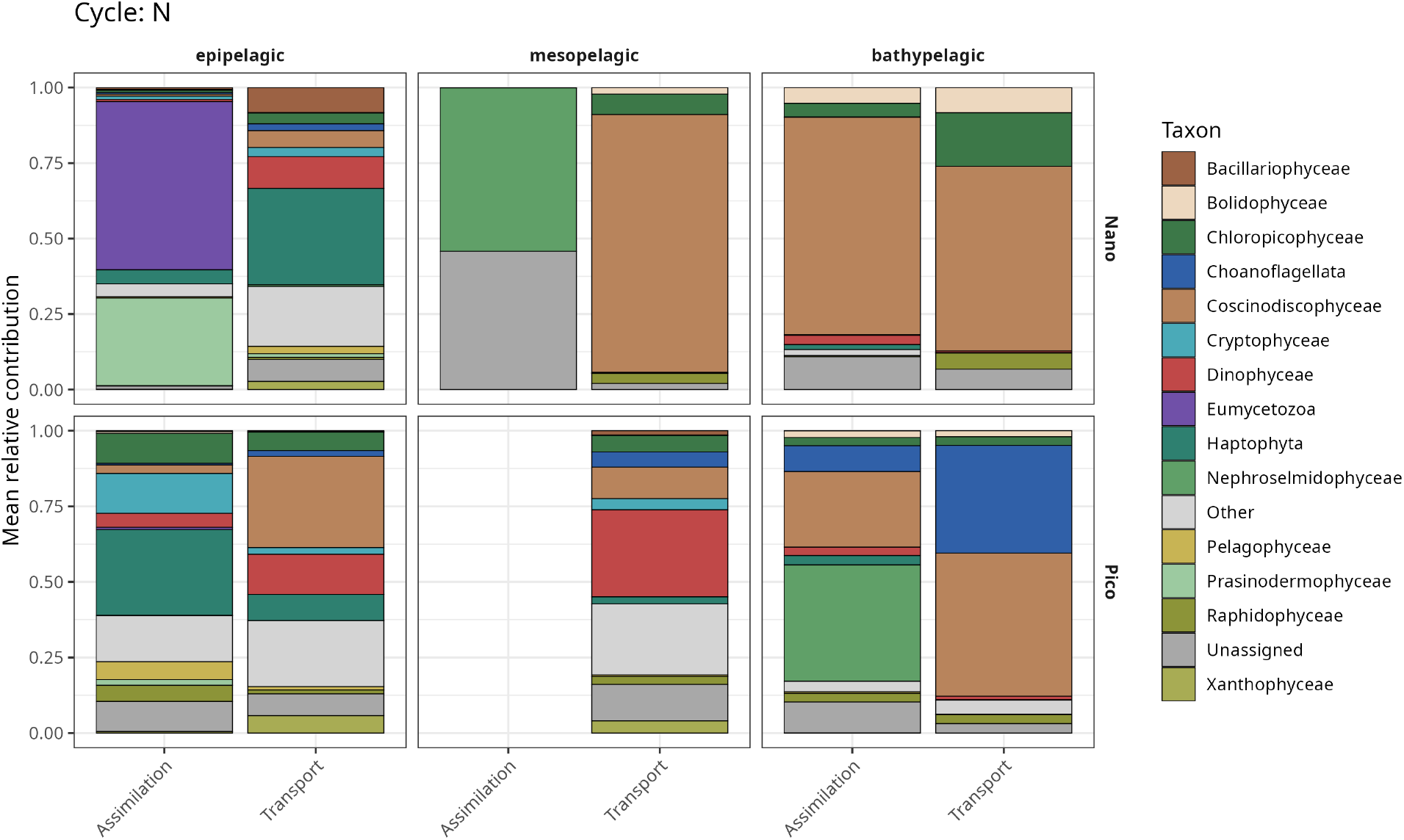
Taxonomic contribution to enriched nitrogen cycling markers. Stacked barplots showing the mean relative contribution of the top eukaryotic taxa to significantly depth-enriched Nitrogen (N) cycling markers, for nano- and pico-size fractions across epipelagic, mesopelagic, and bathypelagic depth zones.

**Fig. S6.**
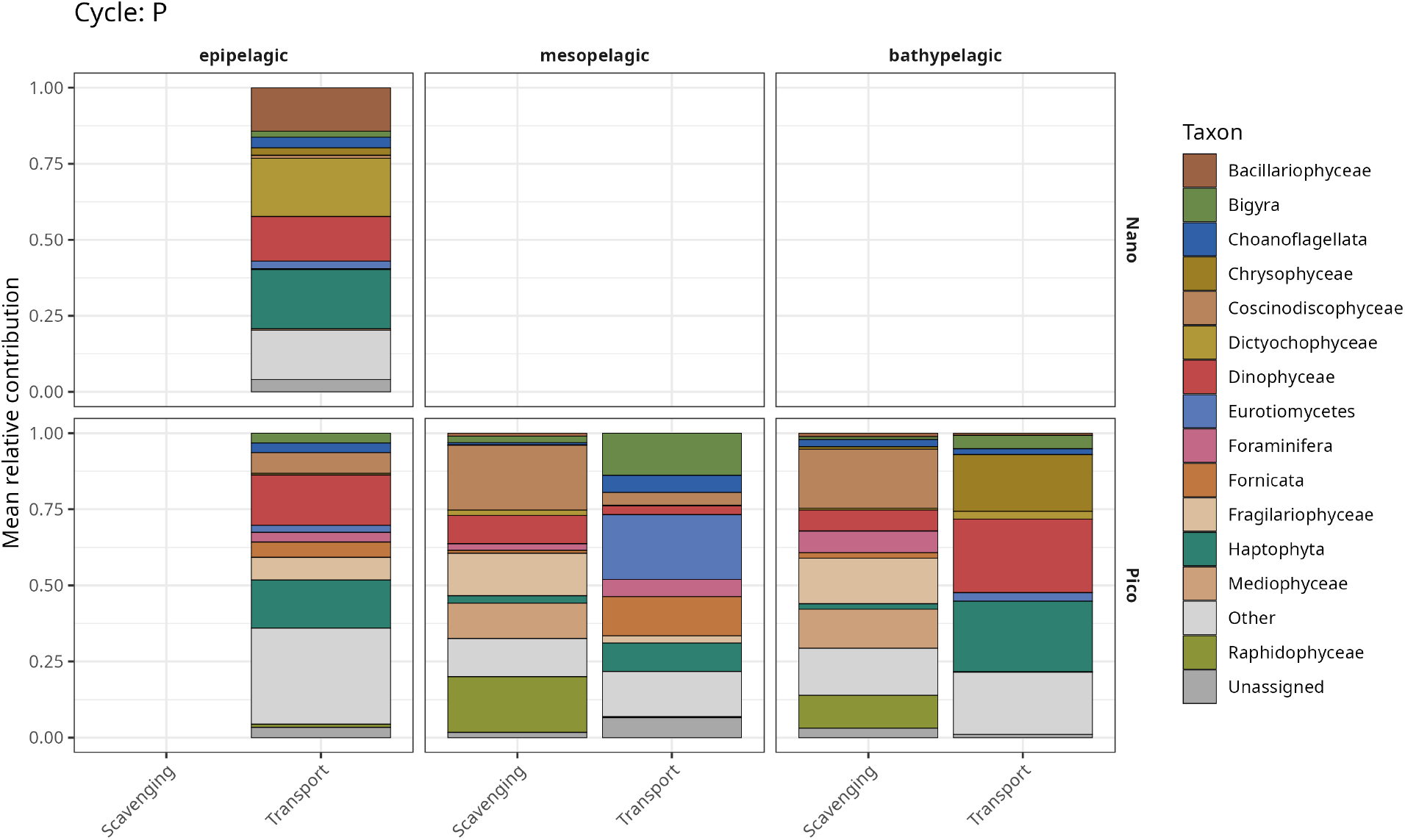
Taxonomic contribution to enriched phosphorus cycling markers. Stacked barplots showing the mean relative contribution of the top eukaryotic taxa to significantly depth-enriched phosphorus (P) cycling markers, for nano- and pico-size fractions across epipelagic, mesopelagic, and bathypelagic depth zones.

**Fig. S7.**
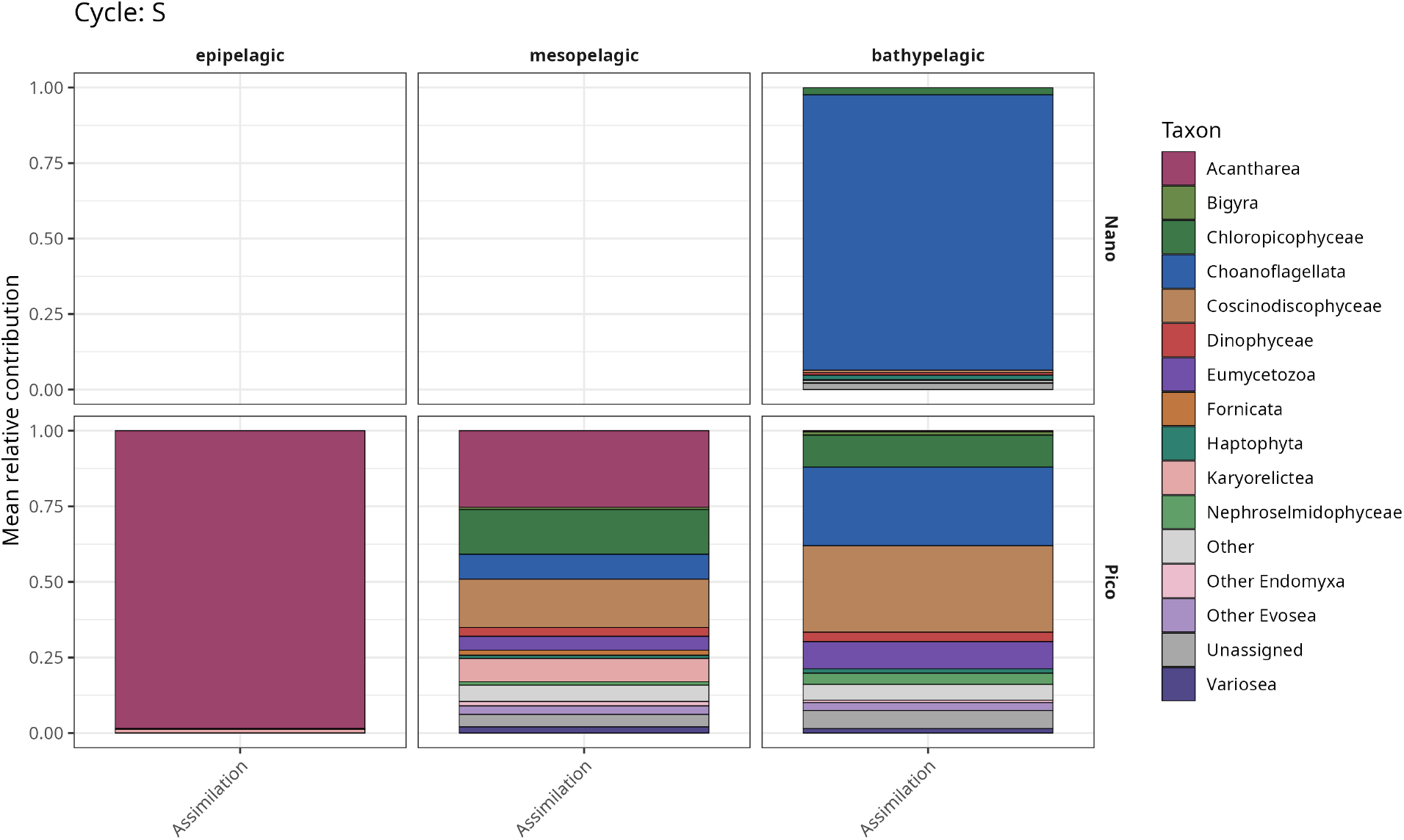
Taxonomic contribution to enriched Sulfur cycling markers. Stacked barplots showing the mean relative contribution of the top eukaryotic taxa to significantly depth-enriched sulfur (S) cycling markers, for nano and pico-size fractions across epipelagic, mesopelagic, and bathypelagic depth zones.

**Fig. S8.**
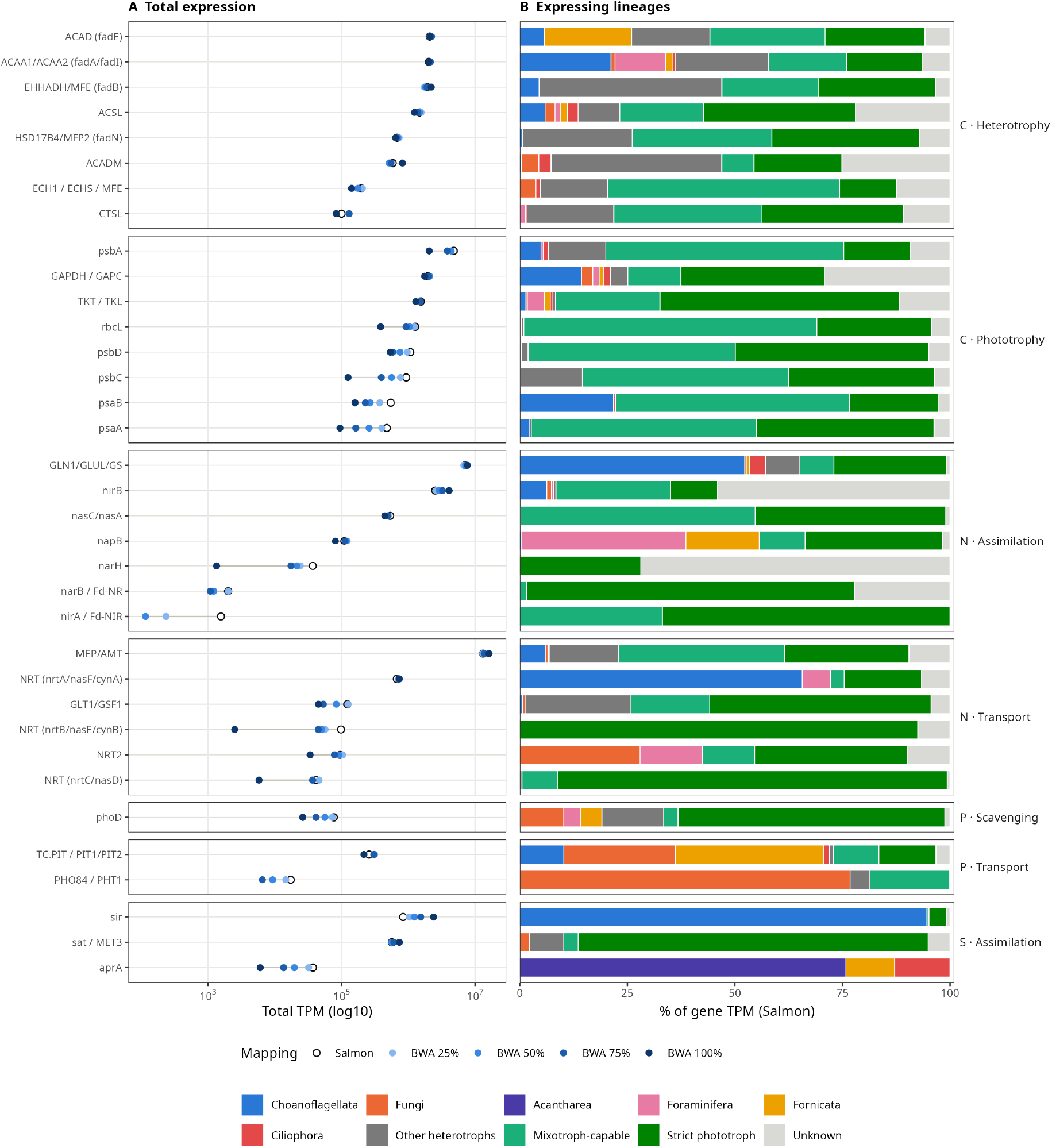
Expression and expressing lineages of eukaryotic metabolic markers in Malaspina bathypelagic metatranscriptomes. Eight most-expressed markers shown for each cycle × functional category, ranked by Salmon TPM; A) Total transcripts per million (TPM, log10 scale) summed across all 51 samples. Open circles show Salmon selective-alignment quantification of metatranscriptomic reads against the catalogue of eukaryotic marker ORFs. Filled blue circles show BWA-MEM2 alignment quantified with CoverM at ≥95% read identity and ≥80% read coverage, using minimum horizontal-coverage thresholds of 25, 50, 75, and 100% of each ORF’s length (light to dark, increasing stringency). Lines connecting the circles for each marker indicate the sensitivity of its signal to quantification method and alignment stringency. (B) Percentage of each marker’s Salmon TPM contributed by each lineage. Trophic categories: obligate heterotrophs, mixotroph-capable taxa, and strict phototrophs; ORFs lacking a resolved phylum are labelled Unknown. Within obligate heterotrophs, Foraminifera, Fornicata, Choanoflagellata, Ciliophora, Fungi, and Acantharea are shown separately, while all other heterotrophic lineages are pooled as “Other heterotrophs.”

**Fig. S9.**
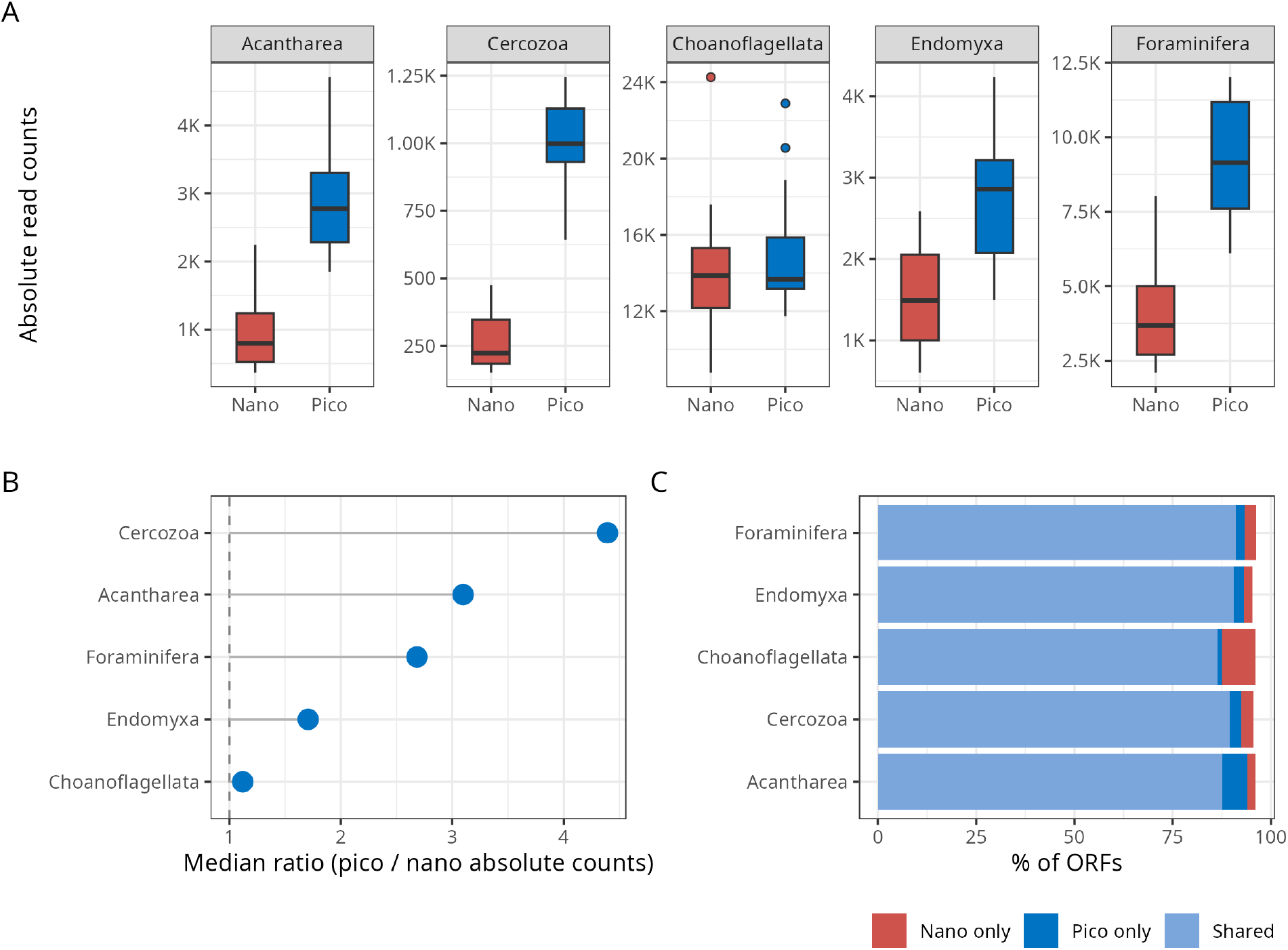
Cross-fraction read counts and ORF overlap for key heterotrophic taxa. (A) Boxplots of absolute read counts (averaged across replicates per station × depth zone) for nano- and pico-size fractions across five target heterotrophic taxa. Only stations with paired nano and pico samples are shown. (B) Median ratio of pico to nano absolute read counts per taxon. A ratio of 1 (dashed line) indicates equal counts across fractions; values below 1 indicate a nano-dominant signal. (C) Percentage of ORFs assigned to each taxon that are detected exclusively in the pico fraction (pico only), exclusively in the nano fraction (nano only), or in both fractions (Shared).

**Fig. S10.**
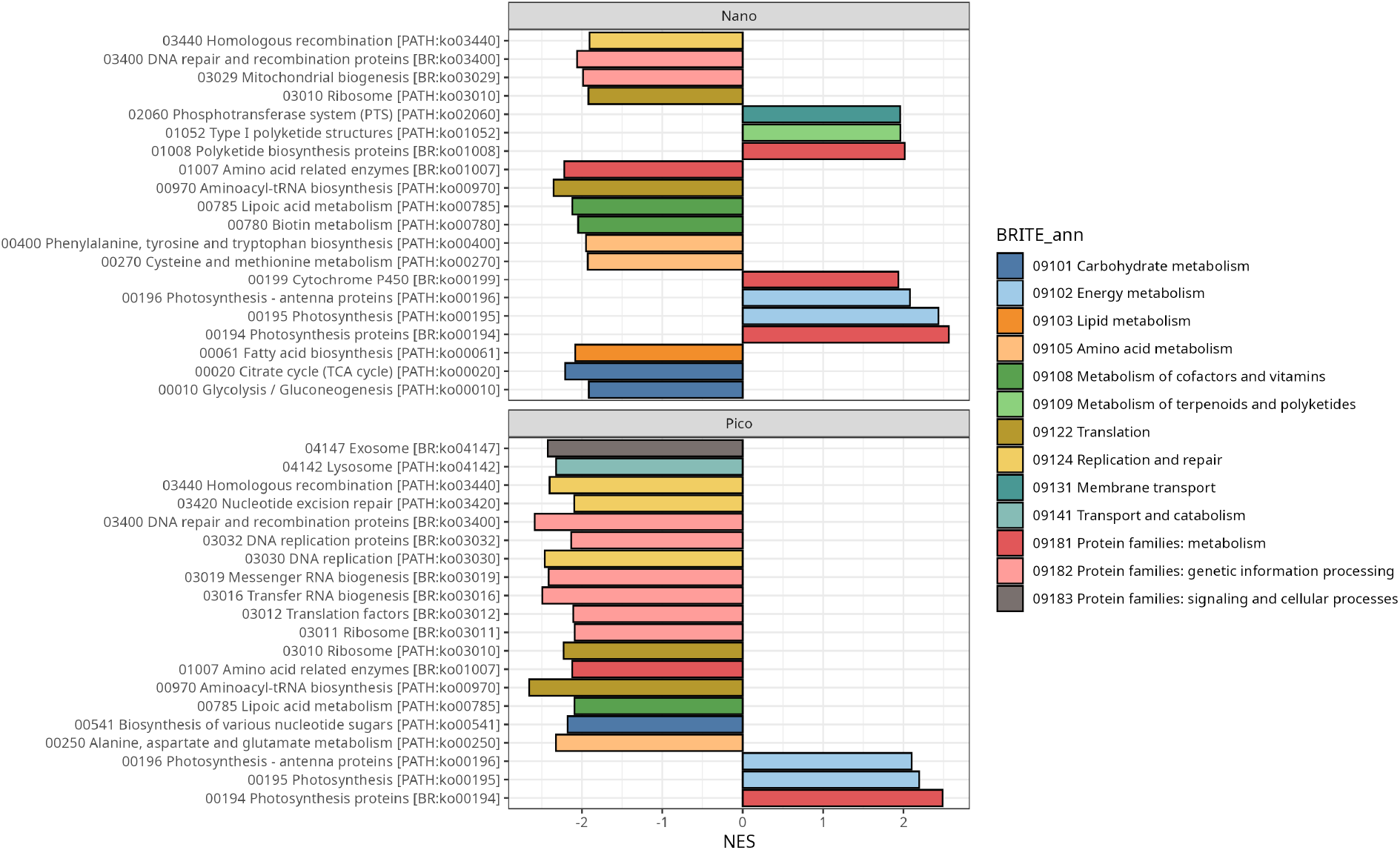
Gene set enrichment analysis of metabolic pathway variability across depth zones. Barplot of normalized enrichment scores (NES) from gene set enrichment analysis (GSEA) of KEGG BRITE functional categories, shown separately for nano and pico eukaryotic size fractions. KOs were ranked by the z-score of their coefficient of variation (CV) in Hellinger-transformed abundance across depth zones. Positive NES values indicate BRITE categories enriched among KOs with high CV (i.e. strongly varying with depth); negative NES values indicate categories enriched among KOs with low CV (i.e. relatively constant across depth zones). Colours denote BRITE functional annotation. Only significantly enriched categories are shown.

**Fig. S11.**
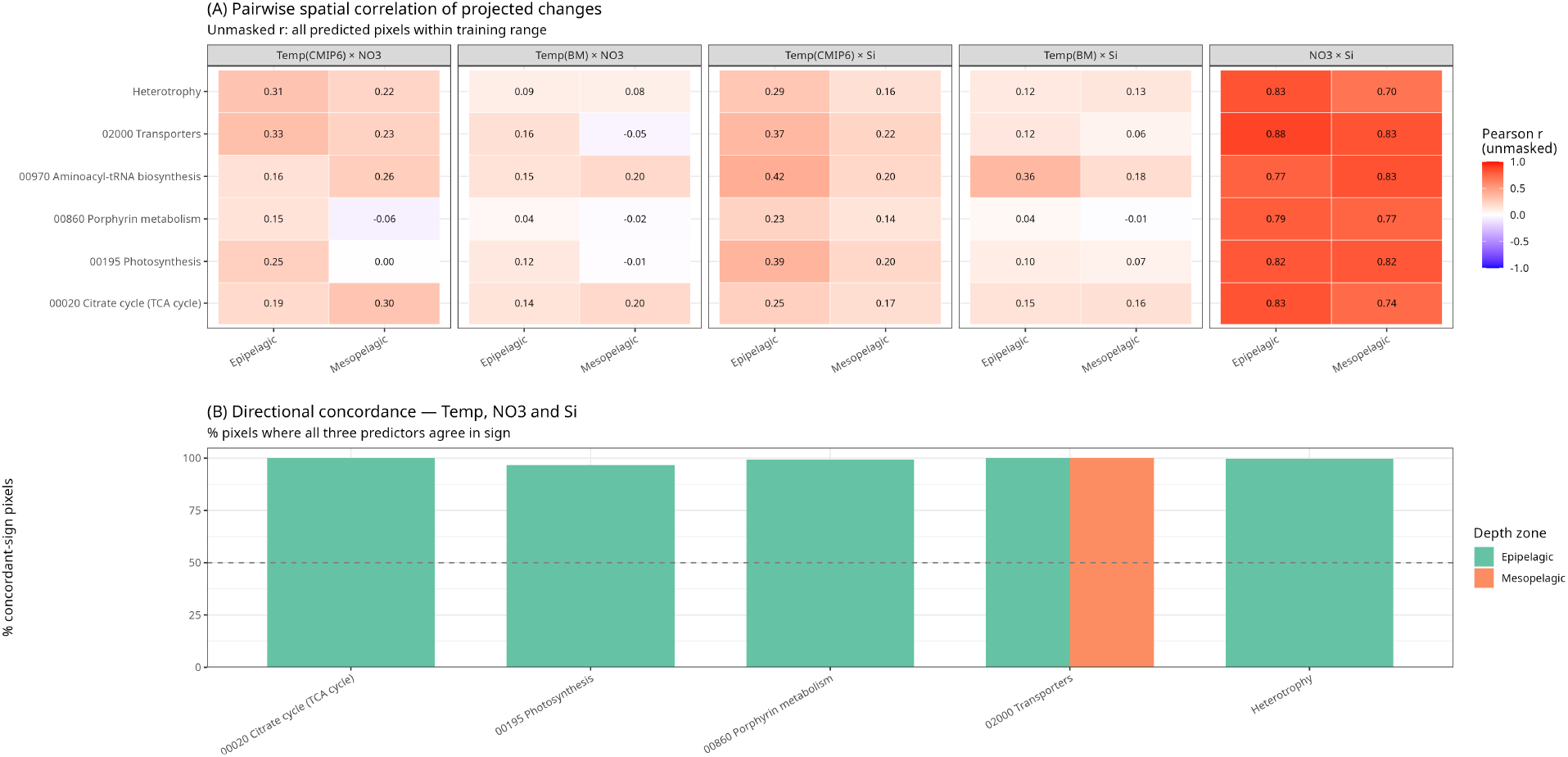
Agreement between temperature- and nutrient-driven projections of picoeukaryote functional potential under SSP5–8.5. (A) Pairwise Pearson correlation between projected percentage-change surfaces for each predictor pair, across all pixels within the training range (unmasked), by functional category and depth zone. Temp(CMIP6) and Temp(BM) denote temperature from the CMIP6 ensemble and from Brito-Morales et al. (2022; the Fig. 5 product), respectively. Masked correlations and full values are in Table S16. (B) Percentage of pixels at which all three predictors (Temp[BM], NO₃, Si) agree in the sign of projected change, among pixels where all three are significant; n above each bar. Aminoacyl-tRNA biosynthesis is omitted (Si yielded no significant pixels). Categories without a mesopelagic bar had no triple-significant pixels there.

**Fig. S12.**
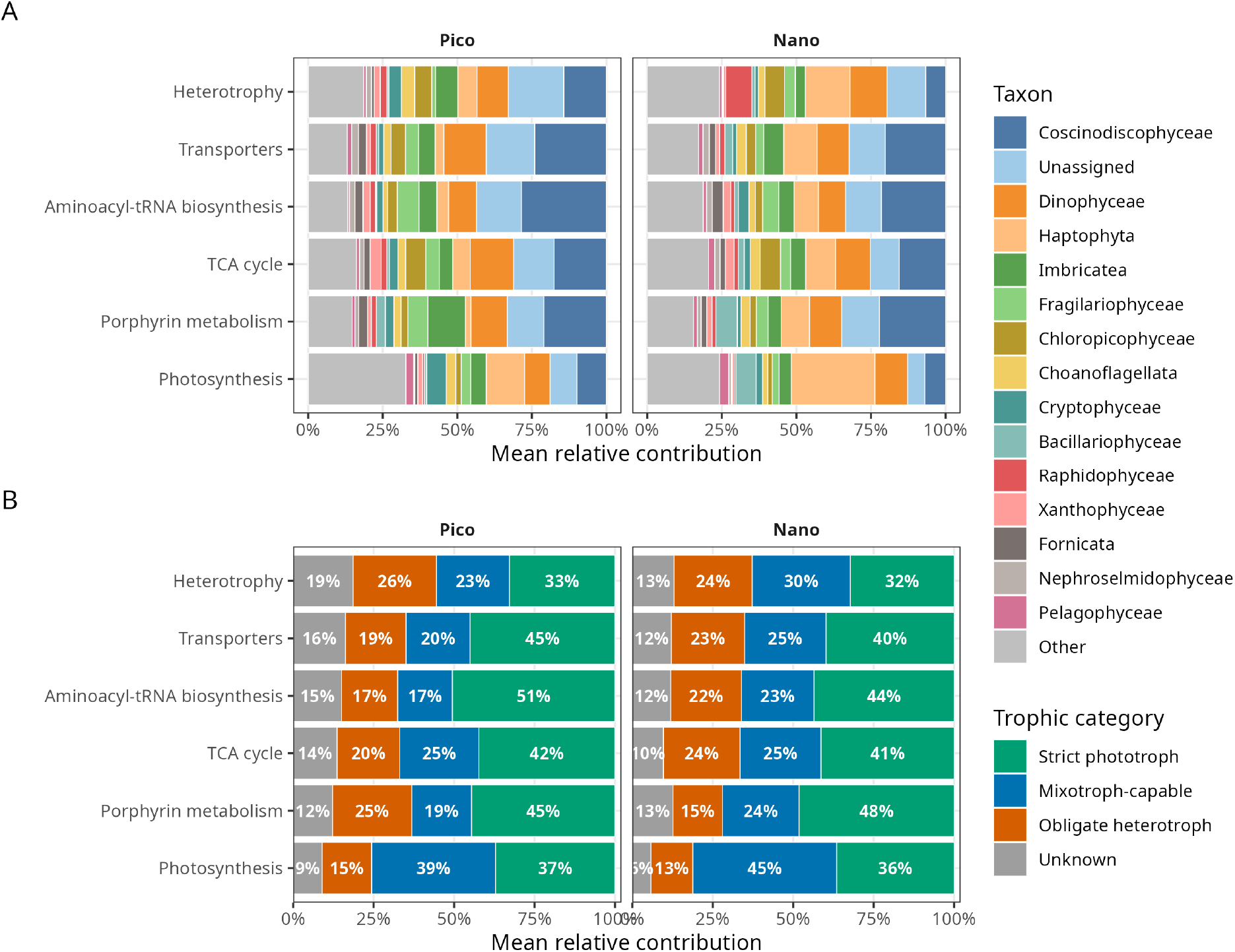
Taxonomic contribution to GAM-modelled functional pathways in Pico (0.2–3 µm) and Nano (3– 20 µm) size fractions. (A) Stacked bar charts showing the mean relative contribution of each taxon (phylum or class when phylum is unknown) to the gene content of each KEGG pathway category considered in the GAM temperature projections. Pathway categories include photosynthesis (00195), TCA cycle (00020), porphyrin and chlorophyll metabolism (00860), aminoacyl-tRNA biosynthesis (00970), transporters (02000), and heterotrophy markers (KOs with significant negative Kendall’s τ correlations with temperature, assigned to the heterotrophy functional class). Contributions are computed as mean per-sample relative abundance (reads assigned to pathway-category genes normalised by total sample read counts), averaged across all samples in each size fraction. The top 15 taxa by total contribution across both fractions are shown individually; remaining taxa are pooled as ‘Other’. (B) The same data aggregated by trophic category. Taxa were assigned to four categories based on the predominant trophic mode at the phylum/class level: strict phototroph (e.g. Bacillariophyta, Chlorophyta, Pelagophyceae, Raphidophyceae, Bolidophyceae); mixotroph-capable (Dinophyceae, Haptophyta, Chrysophyceae, Cryptophyceae and Dictyochophyceae; obligate heterotroph (e.g. Cercozoa, Choanoflagellata, Bigyra, fungi); and unknown (taxa not resolved to a classifiable rank).

## REFERENCES

1. R. Proud, M. J. Cox, A. S. Brierley, Biogeography of the Global Ocean’s Mesopelagic Zone. Current Biology 27, 113–119 (2017).

2. R. Proud, M. Cox, C. Le Guen, A. Brierley, Fine-scale depth structure of pelagic communities throughout the global ocean based on acoustic sound scattering layers. Mar. Ecol. Prog. Ser. 598, 35–48 (2018).

3. P. G. Falkowski, T. Fenchel, E. F. Delong, The microbial engines that drive earth’s biogeochemical cycles. Science 320, 1034–1039 (2008).

4. A. Z. Worden, M. J. Follows, S. J. Giovannoni, S. Wilken, A. E. Zimmerman, P. J. Keeling, Rethinking the marine carbon cycle: Factoring in the multifarious lifestyles of microbes. Science 347, 1257594 (2015).

5. Y. M. Bar-On, R. Milo, The Biomass Composition of the Oceans: A Blueprint of Our Blue Planet. Cell 179, 1451–1454 (2019).

6. D. A. Caron, P. D. Countway, A. C. Jones, D. Y. Kim, A. Schnetzer, Marine Protistan Diversity. Annu. Rev. Mar. Sci. 4, 467–493 (2012).

7. M. Jamy, P. Ramond, D. Bass, J. Del Campo, M. Dunthorn, E. Lara, A. Mitra, D. Vaulot, L. Santoferrara, Towards a trait-based framework for protist ecology and evolution. Trends in Microbiology, S0966842X25002513 (2025).

8. A. Mitra, K. J. Flynn, U. Tillmann, J. A. Raven, D. Caron, D. K. Stoecker, F. Not, P. J. Hansen, G. Hallegraeff, R. Sanders, S. Wilken, G. McManus, M. Johnson, P. Pitta, S. Våge, T. Berge, A. Calbet, F. Thingstad, H. J. Jeong, J. A. Burkholder, P. M. Glibert, E. Granéli, V. Lundgren, Defining Planktonic Protist Functional Groups on Mechanisms for Energy and Nutrient Acquisition: Incorporation of Diverse Mixotrophic Strategies. Protist 167, 106–120 (2016).

9. S. Louca, M. F. Polz, F. Mazel, M. B. N. Albright, J. A. Huber, M. I. O’Connor, M. Ackermann, A. S. Hahn, D. S. Srivastava, S. A. Crowe, M. Doebeli, L. W. Parfrey, Function and functional redundancy in microbial systems. Nature Ecology and Evolution 2, 936–943 (2018).

10. S. Louca, L. Wegener Parfrey, M. Doebeli, Decoupling function and taxonomy in the global ocean microbiome. Science 353, 1272–1277 (2016).

11. P. J. Keeling, Elephants in the room: protists and the importance of morphology and behaviour. Environ Microbiol Rep 5, 1–16 (2013).

12. P. Ramond, M. Sourisseau, N. Simon, S. Romac, S. Schmitt, F. Rigaut-Jalabert, N. Henry, C. de Vargas, R. Siano, Coupling between taxonomic and functional diversity in protistan coastal communities. Environmental Microbiology 21, 730–749 (2019).

13. I. M. Deutschmann, G. Lima-Mendez, A. K. Krabberød, J. Raes, S. M. Vallina, K. Faust, R. Logares, Disentangling environmental effects in microbial association networks. Microbiome 9, 1–18 (2021).

14. C. R. Giner, M. C. Pernice, V. Balagué, C. M. Duarte, J. M. Gasol, R. Logares, R. Massana, Marked changes in diversity and relative activity of picoeukaryotes with depth in the world ocean. ISME Journal 14, 437–449 (2020).

15. P. C. Junger, H. Sarmento, C. R. Giner, M. Mestre, M. Sebastián, X. A. G. Morán, J. Arístegui, S. Agustí, C. M. Duarte, S. G. Acinas, R. Massana, J. M. Gasol, R. Logares, Global biogeography of the smallest plankton across ocean depths. Science Advances 9, eadg9763 (2023).

16. A. Obiol, C. R. Giner, P. Sánchez, C. M. Duarte, S. G. Acinas, R. Massana, A metagenomic assessment of microbial eukaryotic diversity in the global ocean. Molecular Ecology Resources 20, 718–731 (2020).

17. P. Sun, Y. Wang, Y. Zhang, R. Logares, P. Cheng, D. Xu, From the Sunlit to the Aphotic Zone : Assembly Mechanisms and Co-Occurrence Patterns of Protistan-Bacterial Microbiotas in the Western Paci fic Ocean. mSystems (2023).

18. E. Villarino, J. R. Watson, G. Chust, A. J. Woodill, B. Klempay, B. Jonsson, J. M. Gasol, R. Logares, R. Massana, C. R. Giner, G. Salazar, X. A. Alvarez-Salgado, T. S. Catala, C. M. Duarte, S. Agusti, F. Mauro, X. Irigoien, A. D. Barton, Global beta diversity patterns of microbial communities in the surface and deep ocean. Global Ecol Biogeogr 31, 2323–2336 (2022).

19. D. Xu, R. Li, C. Hu, P. Sun, N. Jiao, A. Warren, Microbial Eukaryote Diversity and Activity in the Water Column of the South China Sea Based on DNA and RNA High Throughput Sequencing. Front. Microbiol. 8, 1121 (2017).

20. E. B. Graham, J. E. Knelman, A. Schindlbacher, S. Siciliano, M. Breulmann, A. Yannarell, J. M. Beman, G. Abell, L. Philippot, J. Prosser, A. Foulquier, J. C. Yuste, H. C. Glanville, D. L. Jones, R. Angel, J. Salminen, R. J. Newton, H. Bürgmann, L. J. Ingram, U. Hamer, H. M. P. Siljanen, K. Peltoniemi, K. Potthast, L. Bañeras, M. Hartmann, S. Banerjee, R.-Q. Yu, G. Nogaro, A. Richter, M. Koranda, S. C. Castle, M. Goberna, B. Song, A. Chatterjee, O. C. Nunes, A. R. Lopes, Y. Cao, A. Kaisermann, S. Hallin, M. S. Strickland, J. Garcia-Pausas, J. Barba, H. Kang, K. Isobe, S. Papaspyrou, R. Pastorelli, A. Lagomarsino, E. S. Lindström, N. Basiliko, D. R. Nemergut, Microbes as Engines of Ecosystem Function: When Does Community Structure Enhance Predictions of Ecosystem Processes? Front. Microbiol. 7 (2016).

21. H. P. Grossart, R. Massana, K. D. McMahon, D. A. Walsh, Linking metagenomics to aquatic microbial ecology and biogeochemical cycles. Limnology and Oceanography 65, S2–S20 (2020).

22. Q. Carradec, E. Pelletier, C. Da Silva, A. Alberti, Y. Seeleuthner, R. Blanc-Mathieu, G. Lima-Mendez, F. Rocha, L. Tirichine, K. Labadie, A. Kirilovsky, A. Bertrand, S. Engelen, M.-A. Madoui, R. Méheust, J. Poulain, S. Romac, D. J. Richter, G. Yoshikawa, C. Dimier, S. Kandels-Lewis, M. Picheral, S. Searson, O. Jaillon, J.-M. Aury, E. Karsenti, M. B. Sullivan, S. Sunagawa, P. Bork, F. Not, P. Hingamp, J. Raes, L. Guidi, H. Ogata, C. de Vargas, D. Iudicone, C. Bowler, P. Wincker, A global ocean atlas of eukaryotic genes. Nat Commun 9, 373 (2018).

23. G. Salazar, L. Paoli, A. Alberti, J. Huerta-Cepas, H.-J. Ruscheweyh, M. Cuenca, C. M. Field, L. P. Coelho, C. Cruaud, S. Engelen, A. C. Gregory, K. Labadie, C. Marec, E. Pelletier, M. Royo-Llonch, S. Roux, P. Sánchez, H. Uehara, A. A. Zayed, G. Zeller, M. Carmichael, C. Dimier, J. Ferland, S. Kandels, M. Picheral, S. Pisarev, J. Poulain, S. G. Acinas, M. Babin, P. Bork, E. Boss, C. Bowler, G. Cochrane, C. de Vargas, M. Follows, G. Gorsky, N. Grimsley, L. Guidi, P. Hingamp, D. Iudicone, O. Jaillon, S. Kandels-Lewis, L. Karp-Boss, E. Karsenti, F. Not, H. Ogata, S. Pesant, N. Poulton, J. Raes, C. Sardet, S. Speich, L. Stemmann, M. B. Sullivan, S. Sunagawa, P. Wincker, S. G. Acinas, M. Babin, P. Bork, C. Bowler, C. de Vargas, L. Guidi, P. Hingamp, D. Iudicone, L. Karp-Boss, E. Karsenti, H. Ogata, S. Pesant, S. Speich, M. B. Sullivan, P. Wincker, S. Sunagawa, Gene Expression Changes and Community Turnover Differentially Shape the Global Ocean Metatranscriptome. Cell 179, 1068–1083.e21 (2019).

24. A. J. Perrin, R. G. Dorrell, Protists and protistology in the Anthropocene: challenges for a climate and ecological crisis. BMC Biol 22, 279 (2024).

25. D. A. Hutchins, J. K. Jansson, J. V. Remais, V. I. Rich, B. K. Singh, P. Trivedi, Climate change microbiology — problems and perspectives. Nat Rev Microbiol 17, 391–396 (2019).

26. S. I. Anderson, A. D. Barton, S. Clayton, S. Dutkiewicz, T. A. Rynearson, Marine phytoplankton functional types exhibit diverse responses to thermal change. Nat Commun 12, 6413 (2021).

27. I. Marinov, S. C. Doney, I. D. Lima, Response of ocean phytoplankton community structure to climate change over the 21st century: partitioning the effects of nutrients, temperature and light. Biogeosciences 7, 3941–3959 (2010).

28. A. Tagliabue, ‘Oceans are hugely complex’: modelling marine microbes is key to climate forecasts. Nature 623, 250–252 (2023).

29. C. M. Duarte, Seafaring in the 21st century: The Malaspina 2010 circumnavigation expedition. Limnology and Oceanography Bulletin 24, 11–14 (2015).

30. B. Fox-Kemper, H. T. Hewitt, C. Xiao, G. Aðalgeirsdóttir, S. Drijfhout, T. L. Edwards, N. R. Golledge, M. Hemer, R. E. Kopp, G. Krinner, A. Mix, D. Notz, S. Nowicki, I. S. Nurhati, L. Ruiz, J. Sallée, A. B. A. Slangen, Y. Yu, “Ocean, Cryosphere and Sea Level Change” in Climate Change 2021: The Physical Science Basis. Contribution of Working Group I to the Sixth Assessment Report of the Intergovernmental Panel on Climate Change (Cambridge University Press, ed. 1, 2021; https://www.cambridge.org/core/product/identifier/9781009157896/type/book), pp. 1211–1362.

31. T. Cordier, I. B. Angeles, N. Henry, F. Lejzerowicz, C. Berney, R. Morard, A. Brandt, M.-A. Cambon-Bonavita, L. Guidi, F. Lombard, P. M. Arbizu, R. Massana, C. Orejas, J. Poulain, C. R. Smith, P. Wincker, S. Arnaud-Haond, A. J. Gooday, C. De Vargas, J. Pawlowski, Patterns of eukaryotic diversity from the surface to the deep-ocean sediment. Sci. Adv. 8, eabj9309 (2022).

32. C. de Vargas, S. Audic, N. Henry, J. Decelle, F. Mahé, R. Logares, E. Lara, C. Berney, N. Le Bescot, I. Probert, M. Carmichael, J. Poulain, S. Romac, S. Colin, J.-M. Aury, L. Bittner, S. Chaffron, M. Dunthorn, S. Engelen, O. Flegontova, L. Guidi, A. Horák, O. Jaillon, G. Lima-Mendez, J. Lukeš, S. Malviya, R. Morard, M. Mulot, E. Scalco, R. Siano, F. Vincent, A. Zingone, C. Dimier, M. Picheral, S. Searson, S. Kandels-Lewis, S. G. Acinas, P. Bork, C. Bowler, G. Gorsky, N. Grimsley, P. Hingamp, D. Iudicone, F. Not, H. Ogata, S. Pesant, J. Raes, M. E. Sieracki, S. Speich, L. Stemmann, S. Sunagawa, J. Weissenbach, P. Wincker, E. Karsenti, Ocean plankton. Eukaryotic plankton diversity in the sunlit ocean. Science 348, 1261605 (2015).

33. N. R. Cohen, A. I. Krinos, R. M. Kell, R. J. Chmiel, D. M. Moran, M. R. McIlvin, P. Z. Lopez, A. J. Barth, J. P. Stone, B. A. Alanis, E. W. Chan, J. A. Breier, M. V. Jakuba, R. Johnson, H. Alexander, M. A. Saito, Microeukaryote metabolism across the western North Atlantic Ocean revealed through autonomous underwater profiling. Nat Commun 15, 7325 (2024).

34. S. Agusti, J. I. González-Gordillo, D. Vaqué, M. Estrada, M. I. Cerezo, G. Salazar, J. M. Gasol, C. M. Duarte, Ubiquitous healthy diatoms in the deep sea confirm deep carbon injection by the biological pump. Nat Commun 6, 7608 (2015).

35. S. Chen, Z.-X. Xie, K.-Q. Yan, J.-W. Chen, D.-X. Li, P.-F. Wu, L. Peng, L. Lin, C.-M. Dong, Z. Zhao, G.-Y. Fan, S.-Q. Liu, G. J. Herndl, D.-Z. Wang, Functional vertical connectivity of microbial communities in the ocean. Sci. Adv. 10, eadj8184 (2024).

36. M. Mestre, C. Ruiz-González, R. Logares, C. M. Duarte, J. M. Gasol, M. M. Sala, Sinking particles promote vertical connectivity in the ocean microbiome. Proceedings of the National Academy of Sciences 115, E6799–E6807 (2018).

37. Q. Yang, H. Endo, Y. Yang, A. Ebihara, Y. Yamada, H. Fukuda, T. Nagata, H. Ogata, Metatranscriptomics reveals protistan and viral metabolic reprogramming of sinking particle microbiomes in Oyashio waters. ISME Communications 6, ycag182 (2026).

38. T. M. Huete-Stauffer, R. Logares, M. I. Ansari, A. Røstad, M. L. Calleja, X. A. G. Morán, Increased prokaryotic diversity in the Red Sea deep scattering layer. Environmental Microbiome 18, 87 (2023).

39. P. Ramond, P. E. Galand, R. Logares, Microbial functional diversity and redundancy: moving forward. FEMS Microbiology Reviews 49, fuae031 (2025).

40. R. H. Nilsson, K.-H. Larsson, A. F. S. Taylor, J. Bengtsson-Palme, T. S. Jeppesen, D. Schigel, P. Kennedy, K. Picard, F. O. Glöckner, L. Tedersoo, I. Saar, U. Kõljalg, K. Abarenkov, The UNITE database for molecular identification of fungi: handling dark taxa and parallel taxonomic classifications. Nucleic Acids Res 47, D259–D264 (2019).

41. M. C. Pernice, C. R. Giner, R. Logares, J. Perera-Bel, S. G. Acinas, C. M. Duarte, J. M. Gasol, R. Massana, Large variability of bathypelagic microbial eukaryotic communities across the world’s oceans. The ISME Journal 10, 945–958 (2016).

42. A. B. Bochdansky, M. A. Clouse, G. J. Herndl, Eukaryotic microbes, principally fungi and labyrinthulomycetes, dominate biomass on bathypelagic marine snow. The ISME Journal 11, 362–373 (2017).

43. M. C. Pernice, I. Forn, R. Logares, R. Massana, A fungi hotspot deep in the ocean: explaining the presence of Gjaerumia minor in equatorial Pacific bathypelagic waters. Sci Rep 14, 10601 (2024).

44. E. Breyer, F. Baltar, The largely neglected ecological role of oceanic pelagic fungi. Trends in Ecology & Evolution 38, 870–888 (2023).

45. F. Wang, B. Huang, Y. Xie, S. Cai, X. Wang, J. Mu, Diversity, Composition, and Activities of Nano- and Pico-Eukaryotes in the Northern South China Sea With Influences of Kuroshio Intrusion. Front. Mar. Sci. 8, 658233 (2021).

46. H. Hillebrand, E. Acevedo-Trejos, S. D. Moorthi, A. Ryabov, M. Striebel, P. K. Thomas, M. L. Schneider, Cell size as driver and sentinel of phytoplankton community structure and functioning. Functional Ecology 36, 276–293 (2022).

47. S. T. Dyhrman, B. D. Jenkins, T. A. Rynearson, M. A. Saito, M. L. Mercier, H. Alexander, L. P. Whitney, A. Drzewianowski, V. V. Bulygin, E. M. Bertrand, Z. Wu, C. Benitez-Nelson, A. Heithoff, The Transcriptome and Proteome of the Diatom Thalassiosira pseudonana Reveal a Diverse Phosphorus Stress Response. PLOS ONE 7, e33768 (2012).

48. S. T. Dyhrman, K. C. Ruttenberg, Presence and regulation of alkaline phosphatase activity in eukaryotic phytoplankton from the coastal ocean: Implications for dissolved organic phosphorus remineralization. Limnology and Oceanography 51, 1381–1390 (2006).

49. J. Erez, A. Almogi-Labin, S. Avraham, On the Life History of Planktonic Foraminifera: Lunar Reproduction Cycle in *Globigerinoides Sacculifer* (Brady). Paleoceanography 6, 295–306 (1991).

50. N. Glock, J. Richirt, C. Woehle, C. Algar, M. Armstrong, D. Eichner, H. Firrincieli, A. Makabe, A. Govindankutty Menon, Y. Ishitani, T. Hackl, R. Hubert-Huard, M. Kienast, Y. Milker, A. Mutzberg, S. Ni, S. Okada, S. Rakshit, G. Schmiedl, Z. Steiner, A. Tame, Z. Zhang, H. Nomaki, Widespread occurrence and relevance of phosphate storage in foraminifera. Nature, doi: 10.1038/s41586-024-08431-8 (2025).

51. N. J. Patron, D. G. Durnford, S. Kopriva, Sulfate assimilation in eukaryotes: fusions, relocations and lateral transfers. BMC Evol Biol 8, 39 (2008).

52. P. Sánchez, F. H. Coutinho, M. Sebastián, M. C. Pernice, R. Rodríguez-Martínez, G. Salazar, F. M. Cornejo-Castillo, S. Pesant, X. López-Alforja, E. M. López-García, S. Agustí, T. Gojobori, R. Logares, M. M. Sala, D. Vaqué, R. Massana, C. M. Duarte, S. G. Acinas, J. M. Gasol, Marine picoplankton metagenomes and MAGs from eleven vertical profiles obtained by the Malaspina Expedition. Sci Data 11, 154 (2024).

53. C. A. Durkin, I. Cetinić, M. Estapa, Z. Ljubešić, M. Mucko, A. Neeley, M. Omand, Tracing the path of carbon export in the ocean though DNA sequencing of individual sinking particles. The ISME Journal, 1–11 (2022).

54. P. Tréguer, C. Bowler, B. Moriceau, S. Dutkiewicz, M. Gehlen, O. Aumont, L. Bittner, R. Dugdale, Z. Finkel, D. Iudicone, O. Jahn, L. Guidi, M. Lasbleiz, K. Leblanc, M. Levy, P. Pondaven, Influence of diatom diversity on the ocean biological carbon pump. Nature Geosci 11, 27–37 (2018).

55. J. Arístegui, J. M. Gasol, C. M. Duarte, G. J. Herndld, Microbial oceanography of the dark ocean’s pelagic realm. Limnology & Oceanography 54, 1501–1529 (2009).

56. D. Derilus, M. Z. Rahman, F. Pinero, S. E. Massey, Synergism between the Black Queen effect and the proteomic constraint on genome size reduction in the photosynthetic picoeukaryotes. Sci Rep 10, 8918 (2020).

57. H. L. Kornberg, H. A. Krebs, Synthesis of Cell Constituents from C2-Units by a Modified Tricarboxylic Acid Cycle. Nature 179, 988–991 (1957).

58. M. Nakazawa, T. Minami, K. Teramura, S. Kumamoto, S. Hanato, S. Takenaka, M. Ueda, H. Inui, Y. Nakano, K. Miyatake, Molecular characterization of a bifunctional glyoxylate cycle enzyme, malate synthase/isocitrate lyase, in Euglena gracilis. Comparative Biochemistry and Physiology Part B: Biochemistry and Molecular Biology 141, 445–452 (2005).

59. B. Gašparović, A. Penezić, R. S. Lampitt, N. Sudasinghe, T. Schaub, Free fatty acids, tri-, di- and monoacylglycerol production and depth-related cycling in the Northeast Atlantic. Marine Chemistry 186, 101–109 (2016).

60. M. W. Lomas, S. B. Moran, Evidence for aggregation and export of cyanobacteria and nano-eukaryotes from the Sargasso Sea euphotic zone. Biogeosciences 8, 203–216 (2011).

61. S. Agusti, L. M. Lubián, E. Moreno-Ostos, M. Estrada, C. M. Duarte, Projected Changes in Photosynthetic Picoplankton in a Warmer Subtropical Ocean. Front. Mar. Sci. 5 (2019).

62. M. Cornec, H. Claustre, A. Mignot, L. Guidi, L. Lacour, A. Poteau, F. D’Ortenzio, B. Gentili, C. Schmechtig, Deep Chlorophyll Maxima in the Global Ocean: Occurrences, Drivers and Characteristics. Global Biogeochemical Cycles 35, e2020GB006759 (2021).

63. N. R. Cohen, M. R. McIlvin, D. M. Moran, N. A. Held, J. K. Saunders, N. J. Hawco, M. Brosnahan, G. R. DiTullio, C. Lamborg, J. P. McCrow, C. L. Dupont, A. E. Allen, M. A. Saito, Dinoflagellates alter their carbon and nutrient metabolic strategies across environmental gradients in the central Pacific Ocean. Nat Microbiol 6, 173–186 (2021).

64. S. K. Hu, Z. Liu, H. Alexander, V. Campbell, P. E. Connell, S. T. Dyhrman, K. B. Heidelberg, D. A. Caron, Shifting metabolic priorities among key protistan taxa within and below the euphotic zone. Environmental Microbiology 20, 2865–2879 (2018).

65. D. Padfield, C. Lowe, A. Buckling, R. Ffrench-Constant, Student Research Team, S. Jennings, F. Shelley, J. S. Ólafsson, G. Yvon-Durocher, Metabolic compensation constrains the temperature dependence of gross primary production. Ecology Letters 20, 1250–1260 (2017).

66. Y.-C. Yeh, J. A. Fuhrman, Contrasting diversity patterns of prokaryotes and protists over time and depth at the San-Pedro Ocean Time series. ISME Commun 2, 36 (2022).

67. A. I. Krinos, M. Mars Brisbin, S. K. Hu, N. R. Cohen, T. A. Rynearson, M. J. Follows, F. Schulz, H. Alexander, Missing microbial eukaryotes and misleading meta-omic conclusions. Nat Commun 15, 9873 (2024).

68. P. J. Keeling, F. Burki, H. M. Wilcox, B. Allam, E. E. Allen, L. a. Amaral-Zettler, E. V. Armbrust, J. M. Archibald, A. K. Bharti, C. J. Bell, B. Beszteri, K. D. Bidle, C. T. Cameron, L. Campbell, D. a. Caron, R. A. Cattolico, J. L. Collier, K. Coyne, S. K. Davy, P. Deschamps, S. T. Dyhrman, B. Edvardsen, R. D. Gates, C. J. Gobler, S. J. Greenwood, S. M. Guida, J. L. Jacobi, K. S. Jakobsen, E. R. James, B. Jenkins, U. John, M. D. Johnson, A. R. Juhl, A. Kamp, L. a. Katz, R. Kiene, A. Kudryavtsev, B. S. Leander, S. Lin, C. Lovejoy, D. Lynn, A. Marchetti, G. McManus, A. M. Nedelcu, S. Menden-Deuer, C. Miceli, T. Mock, M. Montresor, M. A. Moran, S. Murray, G. Nadathur, S. Nagai, P. B. Ngam, B. Palenik, J. Pawlowski, G. Petroni, G. Piganeau, M. C. Posewitz, K. Rengefors, G. Romano, M. E. Rumpho, T. Rynearson, K. B. Schilling, D. C. Schroeder, A. G. B. Simpson, C. H. Slamovits, D. R. Smith, G. J. Smith, S. R. Smith, H. M. Sosik, P. Stief, E. Theriot, S. N. Twary, P. E. Umale, D. Vaulot, B. Wawrik, G. L. Wheeler, W. H. Wilson, Y. Xu, A. Zingone, A. Z. Worden, The Marine Microbial Eukaryote Transcriptome Sequencing Project (MMETSP): Illuminating the Functional Diversity of Eukaryotic Life in the Oceans through Transcriptome Sequencing. PLoS Biology 12 (2014).

69. C. D. Prokopowich, T. R. Gregory, T. J. Crease, The correlation between rDNA copy number and genome size in eukaryotes. Genome 46, 48–50 (2003).

70. D. Van Vuuren, B. O’Neill, C. Tebaldi, L. Chini, P. Friedlingstein, T. Hasegawa, K. Riahi, B. Sanderson, B. Govindasamy, N. Bauer, V. Eyring, C. Fall, K. Frieler, M. Gidden, L. Gohar, A. Jones, A. King, R. Knutti, E. Kriegler, P. Lawrence, C. Lennard, J. Lowe, C. Mathison, S. Mehmood, L. Prado, Q. Zhang, S. Rose, A. Ruane, C.-F. Schleussner, R. Seferian, J. Sillmann, C. Smith, A. Sörensson, S. Panickal, K. Tachiiri, N. Vaughan, S. Vishwanathan, T. Yokohata, T. Ziehn, The Scenario Model Intercomparison Project for CMIP7 (ScenarioMIP-CMIP7). Climate and Earth system modeling [Preprint] (2025). 10.5194/egusphere-2024-3765.

71. R. Massana, A. E. Murray, C. M. Preston, E. F. DeLong, Vertical distribution and phylogenetic characterization of marine planktonic Archaea in the Santa Barbara Channel. Appl Environ Microbiol 63, 50–56 (1997).

72. M. Martin, Cutadapt removes adapter sequences from high-throughput sequencing reads. EMBnet.journal 17, 10–12 (2013).

73. D. Li, C. M. Liu, R. Luo, K. Sadakane, T. W. Lam, MEGAHIT: An ultra-fast single-node solution for large and complex metagenomics assembly via succinct de Bruijn graph. Bioinformatics 31, 1674–1676 (2015).

74. P. T. West, A. J. Probst, I. V. Grigoriev, B. C. Thomas, J. F. Banfield, Genome-reconstruction for eukaryotes from complex natural microbial communities. Genome Res. 28, 569–580 (2018).

75. E. L. Karin, M. Mirdita, J. Söding, MetaEuk—sensitive, high-throughput gene discovery, and annotation for large-scale eukaryotic metagenomics. Microbiome 8 (2020).

76. M. Steinegger, J. Söding, Clustering huge protein sequence sets in linear time. Nature Communications 9 (2018).

77. B. Buchfink, C. Xie, D. H. Huson, Fast and sensitive protein alignment using DIAMOND. Nat Methods 12, 59–60 (2015).

78. S. F. Altschul, W. Gish, W. Miller, E. W. Myers, D. J. Lipman, Basic local alignment search tool. Journal of Molecular Biology 215, 403–410 (1990).

79. M. Kanehisa, S. Gotu, KEGG: Kyoto Encyclopedia of Genes and Genomes. Nucleic Acids Research 28, 27–30 (2000).

80. M. Steinegger, J. Söding, MMseqs2 enables sensitive protein sequence searching for the analysis of massive data sets. Nat Biotechnol 35, 1026–1028 (2017).

81. D. J. Richter, C. Berney, J. F. H. Strassert, F. Burki, C. de Vargas, EukProt: A database of genome-scale predicted proteins across the diversity of eukaryotic life. Peer Community Journal, 1–11 (2020).

82. R. D. Groussman, S. Blaskowski, S. N. Coesel, E. V. Armbrust, MarFERReT, an open-source, version-controlled reference library of marine microbial eukaryote functional genes. Sci Data 10, 926 (2023).

83. M. Mirdita, L. von den Driesch, C. Galiez, M. J. Martin, J. Söding, M. Steinegger, Uniclust databases of clustered and deeply annotated protein sequences and alignments. Nucleic Acids Res 45, D170–D176 (2017).

84. S. A. Chamberlain, E. Szöcs, taxize: taxonomic search and retrieval in R. F1000Res 2, 191 (2013).

85. European Union-Copernicus Marine Service, Global ocean biogeochemistry hindcast, Mercator Ocean International (2018); 10.48670/MOI-00019.

86. R Core Team, R: A Language and Environment for Statistical Computing (R Foundation for Statistical Computing, Vienna, Austria, 2023; https://www.R-project.org/).

87. A. Kassambara, ggpubr: “ggplot2” Based Publication Ready Plots, (2016); 10.32614/CRAN.package.ggpubr.

88. N. J. Freyria, T. C. De Oliveira, A. Meng, E. Pelletier, C. Lovejoy, Shotgun metagenomics reveals the flexibility and diversity of Arctic marine microbiomes. ISME Communications 5, ycaf007 (2025).

89. S. J. Gleich, S. K. Hu, A. I. Krinos, D. A. Caron, Protistan community composition and metabolism in the North Pacific Subtropical Gyre: Influences of mesoscale eddies and depth. Environmental Microbiology 26, e16556 (2024).

90. S. González-Motos, L. Montiel, V. Balagué, F.-Y. Bouget, R. Massana, J. M. Gasol, J. M. González, P. E. Galand, R. Logares, Functional redundancy enables emergent metabolic dynamics in marine microbiomes. bioRxiv [Preprint] (2025). 10.1101/2025.08.12.669827.

91. T. Wu, E. Hu, S. Xu, M. Chen, P. Guo, Z. Dai, T. Feng, L. Zhou, W. Tang, L. Zhan, X. Fu, S. Liu, X. Bo, G. Yu, clusterProfiler 4.0: A universal enrichment tool for interpreting omics data. The Innovation 2, 100141 (2021).

92. I. Sanz-Sáez, A. G. Bravo, M. Ferri, J.-M. Carreras, O. Sánchez, M. Sebastian, C. Ruiz-González, E. Capo, C. M. Duarte, J. M. Gasol, P. Sánchez, S. G. Acinas, Microorganisms Involved in Methylmercury Demethylation and Mercury Reduction are Widely Distributed and Active in the Bathypelagic Deep Ocean Waters. Environ. Sci. Technol. 58, 13795–13807 (2024).

93. R. Patro, G. Duggal, M. I. Love, R. A. Irizarry, C. Kingsford, Salmon provides fast and bias-aware quantification of transcript expression. Nat Methods 14, 417–419 (2017).

94. Md. Vasimuddin, S. Misra, H. Li, S. Aluru, “Efficient Architecture-Aware Acceleration of BWA-MEM for Multicore Systems” in 2019 IEEE International Parallel and Distributed Processing Symposium (IPDPS) (IEEE, Rio de Janeiro, Brazil, 2019; https://ieeexplore.ieee.org/document/8820962/), pp. 314–324.

95. S. T. N. Aroney, R. J. P. Newell, J. N. Nissen, A. P. Camargo, G. W. Tyson, B. J. Woodcroft, CoverM: read alignment statistics for metagenomics. Bioinformatics 41, btaf147 (2025).

96. S. N. Wood, Generalized Additive Models: An Introduction with R (Chapman and Hall/CRC, ed. 2, 2017; https://www.taylorfrancis.com/books/9781498728348).

97. I. Brito-Morales, D. S. Schoeman, J. D. Everett, C. J. Klein, D. C. Dunn, J. García Molinos, M. T. Burrows, K. C. V. Buenafe, R. M. Dominguez, H. P. Possingham, A. J. Richardson, Towards climate-smart, three-dimensional protected areas for biodiversity conservation in the high seas. Nat. Clim. Chang. 12, 402–407 (2022).\

